# Red-shifted GRAB acetylcholine sensors for multiplex imaging *in vivo*

**DOI:** 10.1101/2024.12.22.627112

**Authors:** Shu Xie, Xiaolei Miao, Guochuan Li, Yu Zheng, Mengyao Li, En Ji, Jinxu Wang, Shaochuang Li, Ruyi Cai, Lan Geng, Jiesi Feng, Changwei Wei, Yulong Li

## Abstract

The neurotransmitter acetylcholine (ACh) is essential in both the central and peripheral nervous systems. Recent studies highlight the significance of interactions between ACh and various neuromodulators in regulating complex behaviors. The ability to simultaneously image ACh and other neuromodulators can provide valuable information regarding the mechanisms underlying these behaviors. Here, we developed a series of red fluorescent G protein-coupled receptor activation-based (GRAB) ACh sensors, with a wide detection range and expanded spectral profile. The high-affinity sensor, rACh1h, reliably detects ACh release in various brain regions, including the nucleus accumbens, amygdala, hippocampus, and cortex. Moreover, rACh1h can be co-expressed with green fluorescent sensors in order to record ACh release together with other neurochemicals in various behavioral contexts using fiber photometry and two-photon imaging, with high spatiotemporal resolution. These new ACh sensors can therefore provide valuable new insights regarding the functional role of the cholinergic system under both physiological and pathological conditions.

## Main

Acetylcholine (ACh), the first neurotransmitter to be identified, plays important roles in both the central and peripheral nervous systems^1,2,3,4,5,6^. In the brain, cholinergic neurons are involved in diverse functions, including attention, arousal, associative learning, and regulating the sleep-wake cycle^6,7^. In addition, the crosstalk between ACh and other neurochemicals has been reported to mediate motivation, cue detection, and reinforcement learning^8,9^. In the striatum, the release of ACh can be inhibited by dopamine (DA) through dopamine D2 receptors^10,11^, while it is driven by glutamine (Glu) inputs from cortical thalalmus^12^. This modulation of ACh by dopaminergic and glutaminergic internation is essential for decision making and learning processes. Pioneer research also indicated that the interaction between ACh and oxytocin in the hippocampus is crucial for regulating brain stages^13^. The simultaneously imaging ACh and other neurochemicals has provided valuable insights into the regulation of brain functions controlled by these signaling processes^14^. Such investigations are helpful in identifying new drug targets and developing innovative therapeutic strategies for neural diseases^15^. Current state-of-the-art green ACh sensors such as GRAB_ACh3.0_ and iAChSnFR are based on green fluorescent proteins and have been used to measure ACh *in vivo*^16–18;^ however, a red fluorescent ACh sensor would be extremely valuable due to its spectral compatibility with green fluorescent sensors, allowing for the simultaneous detection of ACh and other neurochemicals.

Here, we developed a series of red fluorescent ACh sensors. These red-shifted sensors, which we call rACh1h, rACh1m, and rACh1l (with high, medium, and low ACh affinity, respectively), have a >500% increase in fluorescence in response to ACh. We then compared the performance—including the response to ACh and the signal-to-noise ratio (SNR)—of these red fluorescent ACh sensors with GRAB_ACh3.0_ and iAChSnFR. We also show that rACh1h can be used to monitor both spontaneous and optogenetically evoked endogenous ACh release *in vivo* using fiber photometry. When coupled with green GRAB sensors in dual-color recordings, rACh1h revealed a strong correlation between ACh and DA signals in Pavlovian conditioning tasks, as well as distinct dynamics of ACh and serotonin (5-HT) in sleep-wake cycles. Furthermore, multiplex imaging using two-photon microscopy elucidated the release patterns of ACh and norepinephrine (NE) across various behaviors in the visual cortex. Thus, these red-shifted indicators provide a new toolkit for investigating the functional roles of ACh in both health and diseases.

## Results

### Development and characterization of red ACh sensors

To expand the spectral profile of GRAB ACh sensors, we generated a series of red fluorescent ACh sensors. We began by transplanting the cpmApple module from the red fluorescent dopamine sensor rGRAB_DA_ into the third intracellular loop of the mouse type 3 muscarinic ACh receptor (M_3_R) (Fig. 1a), followed by systematic optimization of both the receptor and the fluorescent module (Extended Data Fig. 1)^19–21^. Screening approximately 2000 variants using the ACh-induced change in fluorescence led to the low-affinity rACh1l ACh sensor. Given that the green fluorescent GRAB sensor ACh3.0, which is based on human M_3_R, produces a large change in fluorescence upon binding ACh^17^, we attempted to improve the response and affinity of the red fluorescent sensor using a chimeric strategy in which we fused the sequence of human M_3_R before the 4.55 site with rACh1l after the 4.55 site^20^. Screening >1,000 candidates using the ACh-induced change in fluorescence and the affinity index (see Methods for details), we obtained a high-affinity rACh1h sensor, which has an improved response and higher affinity compared to rACh1l. We then generated a medium-affinity sensor, rACh1m, by introducing the N513^6.58^ K substitution in rACh1h. Finally, we introduced the W199^4.57^ T mutation in rACh1h in order to create an ACh-insensitive version, rAChmut, to serve as a negative control.

**Fig. 1.**
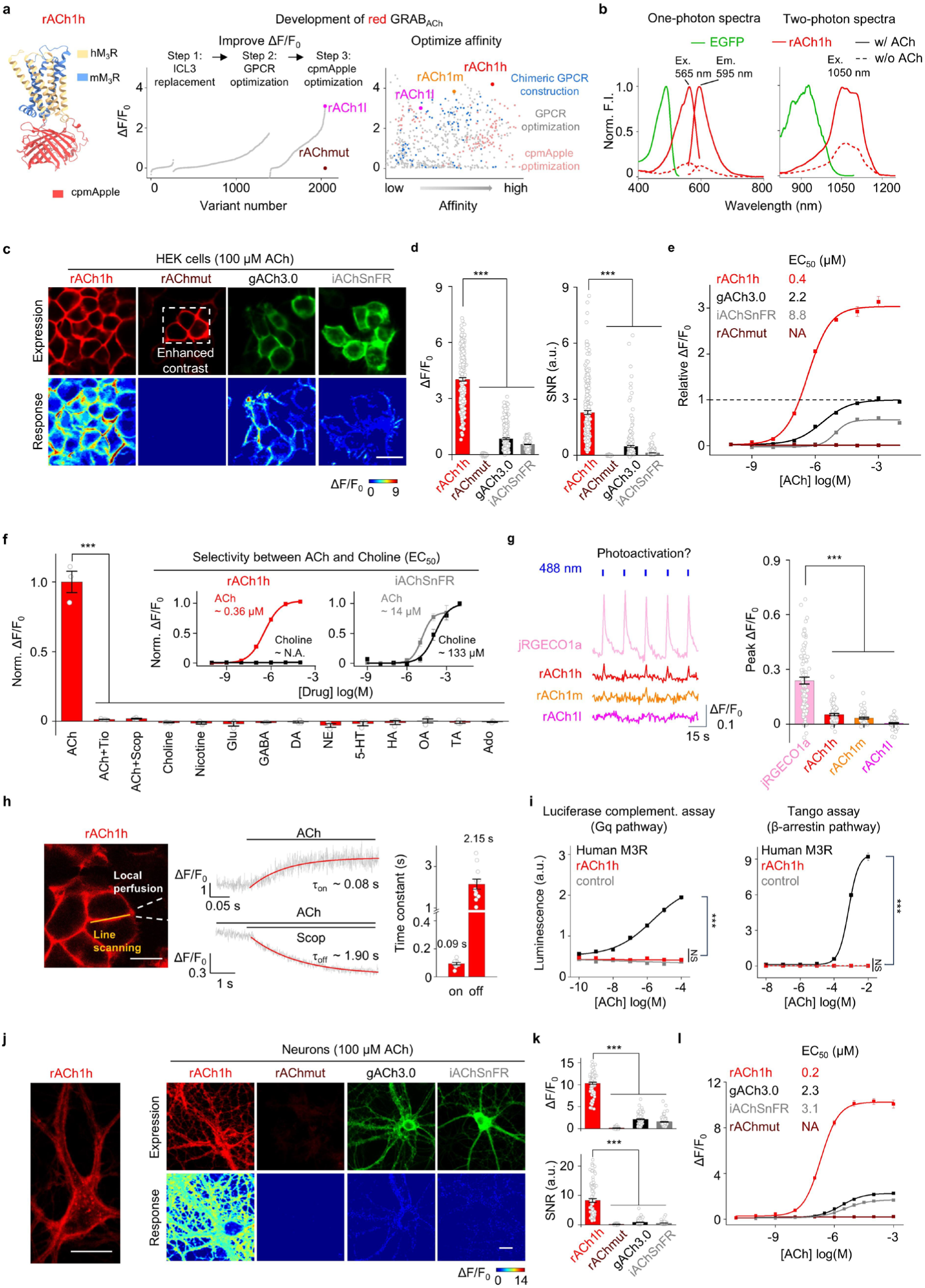
Development and performance of red GRAB_ACh_ sensors. **a**, The predicted structure generated using AlphaFold^42^ (left) and development (right) of red ACh sensors. **b**, One photon and two photon spectral profiles of rACh1h in the absence (dashed line) or presence of 100 μM ACh (solid line). Left, one photon excitation and emission spectra. Right, two photon spectra. Data of EGFP is acquired from FPbase^43^. **c**, Representative images showing expression (top) and response (bottom) to 100 μM ACh of rACh1h, rAChmut, gACh3.0 and iAChSnFR. Scale bar, 20 μm. **d**, Quantification of ΔF/F_0_ and signal-to-noise ratio of rACh1h, rAChmut, gACh3.0 and iAChSnFR before and after 100 μM ACh addition, mean ± s.e.m. n = 240/6, 182/5, 245/7 and 215/6 for rACh1h, rAChmut, gACh3.0 and iAChSnFR respectively. One-way ANOVA with post hoc Tukey’s test was performed. Post hoc test: for ΔF/F_0_ and SNR, P = 0 for rACh1h versus other sensors. **e**, Normalized dose-response curves of rACh1h, rAChmut, gACh3.0 and iAChSnFR. n = 3 wells for each sensor, with 300 - 500 cells per well. **f**, Pharmacological specificity of rACh1h in HEK cells. Tiotropium (Tio), M3R antagonist; Scopolamine (Scop), M3R antagonist; Glu, glutamate; GABA, γ-aminobutyric acid; DA, dopamine; NE, norepinephrine; 5-HT, serotonin; HA, histamine; OA, octopamine; TA, tyramine; Ado, adenosine. Antagonists were applied at 100 μM, others at 10 μM. n = 3 wells for rACh1h,300 - 500 cells per well, mean ± s.e.m. One-way ANOVA with post hoc Tukey’s test was performed, post hoc test: P = 0 for ACh versus ACh + Tio, ACh + Scop and other compounds. The insets show dose–response curves for ACh and choline; n = 3 wells with 300 - 500 cells per well each. **g**, Representative traces (Left) and peak ΔF/F_0_ (Right) in response to blue light in cells expressing jRGECO1a and rACh1h, n = 82/5 and 73/5 for jRGECO1a and rACh1h. Two-tailed Student’s t-test was performed, P = 6.4 × 10^-14^ between jRGECO1a and rACh1h. **h**, Kinetics measurements of rACh1h. Schematic illustration showing the experimental setup of line-scanning and local puffing, Scale bar, 20 μm. (Left), representative traces of sensor fluorescence increase in response to ACh (Medium top) and decrease in response to Scop (Medium bottom). Group summary of on and off kinetics for the sensors (Right), mean ± s.e.m. n = 9/4 for rACh1h on kinetics; n = 11/3 for rACh1h off kinetics. **i**, Downstream coupling test. Human M3R; rACh1h; Control, without expression of WT M3R or sensors. For the luciferase complementation assay, n = 3 wells per group, One-way ANOVA with post hoc Tukey’s test was performed, post hoc test: P = 0 for rACh1h versus Human M3R. For the tango assay, n = 3 wells per group, ANOVA with post hoc Tukey’s test was performed, post hoc test: P = 0 and 1 for rACh1h versus Human M3R and control respectively. **j**, Representative images (Left) of cultured neurons expressing rACh1h. Scale bar, 20 μm. Expression and response (Right) of rACh1h, rAChmut, gACh3.0 and iAChSnFR in cultured neurons with 100 μM ACh addition. **k**, Group summary of ΔF/F_0_ and SNR of rACh1h, rAChmut, gACh3.0 and iAChSnFR. n = 90/3 for each sensors, mean ± s.e.m. One-way ANOVA with post hoc Tukey’s test was performed. For ΔF/F_0_, post hoc test: P = 0 for rACh1h versus others. For SNR, post hoc test: P = 0 for rACh1h versus gACh3.0 and iAChSnFR; P = 5.26×10^-8^ between rACh1h and rAChmut. **l**, Dose-response curves for ACh sensors. n = 90/3 for each sensor. EC_50_, half-maximum effective concentration; FI, fluorescence intensity; NA, not available; NS, not significant.

We then expressed these rACh sensors in HEK293T cells and characterized their spectral properties using both one-photon and two-photon excitation. We found that rACh1h has excitation peaks at 565 nm (one-photon) and 1050 nm (two-photon) (Fig.1b). Similarly, rACh1m and rACh1l exhibit one-photon excitation peaks at 560 nm, with two-photon excitation peaks at 1060 nm and 1110 nm, respectively (Extended Data Fig.2a-b). The red fluorescent ACh sensors have a robust increase in fluorescence (ΔF/F_0_) in response to 100 μM ACh, with rACh1h having a peak ΔF/F_0_ of approximately 500%; in contrast, ACh has no effect when applied to cells expressing rAChmut (Fig. 1c-d and Extended Data Fig. 2c-d). Importantly, rACh1h has a higher response to ACh, with a higher SNR, than both gACh3.0 and iAChSnFR. Specifically, dose-response curves showed that rACh1h has a half-maximum effective concentration (EC_50_) of ∼0.4 μM compared to 2.2 μM and 8.8 μM for gACh3.0 and iAChSnFR, respectively (Fig. 1e). Moreover, rACh1m and rACh1l have EC_50_ values of ∼1.2 μM and 4 μM, respectively (Extended Data Fig. 2e).

After being released from the presynaptic terminal, ACh is degraded to choline by acetylcholinesterase in the synaptic cleft^22^. We therefore measured the selectivity of ACh sensors for ACh over choline and found that our red fluorescent ACh sensors inherited the parent receptor’s pharmacological specificity and had no detectable response to choline, while iAChSnFR responded to both ACh and choline (Fig. 1f and Extended Data Fig. 2f). Furthermore, the red fluorescent ACh sensors did not respond to any other signaling molecules tested, including a wide variety of neurotransmitters and neuromodulators (Fig. 1f and Extended Data Fig. 2g). Previous studies found that cpmApple-based sensors can be photoactivated by blue light^23,24^; however, we found that blue (488-nm) light elicited only a small increase in fluorescence (with ΔF/F_0_ values of ∼5%, ∼3%, and ∼0.5% for rACh1h, rACh1m, and rACh1l, respectively), compared to a ∼25% increase in jRGECO1a fluorescence (Fig. 1g and Extended Data Fig. 2h-i).

To measure the kinetics of our red fluorescent ACh sensors, we expressed them in HEK293T cells and performed rapid line-scanning microscopy while applying a local puff of ACh (to measure the activation time constant, *τ*_on_), followed by the ACh receptor antagonist scopolamine (to measure *τ*_off_) (Fig. 1h and Extended Data Fig. 3a-c). Our analysis revealed a *τ*_on_ value of approximately 0.1s for all three ACh sensors, and *τ*_off_ values ranging from 1.37 s to 2.15 s, reflecting the sensors’ differences in affinity.

To confirm that the ACh sensors do not couple to downstream signaling pathways, we used the luciferase complementation assay^25^ and the Tango assay^26^ to measure the G protein and β-arrestin pathways, respectively. As expected, wild-type human M_3_R exhibited robust coupling, while none of the three red fluorescent ACh sensors had measurable coupling (Fig. 1i and Extended Data Fig. 3d). Importantly, the ACh-induced increase in fluorescence was stable for at least 2 hours, with minimal arrestin-mediated internalization (Extended Data Fig. 3e), indicating that these sensors can be used for long-term monitoring of ACh dynamics.

Next, we tested the performance of our ACh sensors in cultured cortical neurons. Consistent with our results obtained with HEK293T cells, all of the red fluorescent sensors were expressed at robust levels in the plasma membrane (Fig. 1j and Extended Data Fig. 3f). Moreover, upon application of 100 μM ACh, rACh1h, rACh1m, and rACh1l exhibited a fluorescence increase of ∼1000%, 800%, and 680%, respectively, while the ACh-insensitive rAChmut sensor had no detectable response (Fig. 1k and Extended Data Fig. 3g). In addition, rACh1h had a significantly higher fluorescence response and a higher SNR compared to both gACh3.0 and iAChSnFR. Dose-response curves measured in cultured neurons revealed EC_50_ values of ∼0.2 μM, 0.5 μM, and 2.8 μM for rACh1h, rACh1m, and rACh1l, respectively (Fig. 1l and Extended Data Fig. 3h). Finally, and consistent with our findings in HEK283T cells, rACh1h had higher affinity compared to both gACh3.0 and iAChSnFR. Together, these data suggest that our red fluorescent ACh sensors are suitable for use in cultured neurons, and rACh1h outperforms existing green fluorescent ACh sensors in terms of the response, SNR, and ligand affinity.

### Detecting ACh dynamics in acute brain slices

Prior studies showed that ACh plays an important functional role in the striatum^8,11,27^. To test whether our red-shifted ACh sensors can report the release of endogenous ACh, we injected an adeno-associated virus (AAV) expressing the rACh1h sensor into the nucleus accumbens (NAc), a structure that contains cholinergic interneurons. Three weeks after virus injection, we prepared acute brain slices and performed two-photon imaging while applying electrical stimuli to induce ACh release (Fig. 2a). We positioned the stimulating electrode in the NAc and applied increasing numbers of electrical pulses (delivered at 20 Hz) to the brain slice (Fig. 2b). We measured a stimulus number–dependent increase in rACh1h fluorescence, with 100 pulses producing an increase of ∼20%; moreover, the response was significantly inhibited by the M_3_R antagonist scop (Fig. 2c-d). We then measured the kinetics of the change in rACh1h fluorescent in response to a single electrical pulse (Fig.2e), with *τ*_on_ and *τ*_off_ values of ∼0.08 s and 3.7 s, respectively (Fig. 2f). These results indicate that the rACh1h sensor can reliably detect the release of endogenous ACh in acute brain slices.

**Fig. 2.**
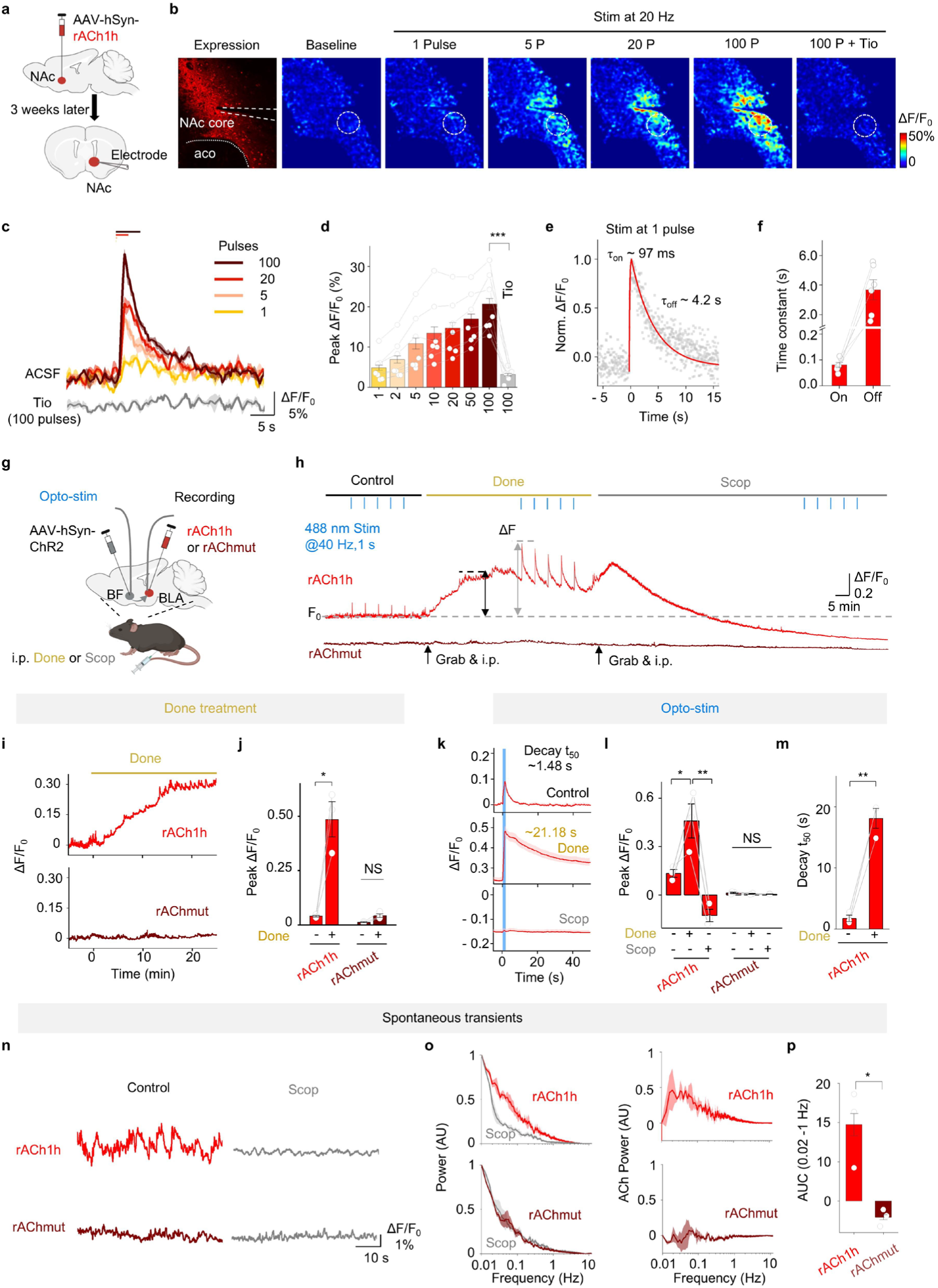
Detection of ACh dynamics *ex vivo* and *in vivo*. **a**, Schematic illustration depicting the two-photon imaging of acute brain slices prepared from mice expressing rACh1h in the NAc. An electrode placed in the NAc was used to evoke release of endogenous ACh. **b**, Representative images of expression of rACh1h response to electrical evoked ACh. The dashed circles indicate the ROI used to calculate the response, and the approximate location of the stimulating electrode is indicated. Scale bar, 100 μm. **c**, Representative traces of the fluorescence change in rACh1h to electrical stimulation. **d**, Group summary of the fluorescence change in rACh1h to electrical stimulation. mean ± s.e.m. n = 7 slices from 3 mice. Two-tailed Student’s t tests: P = 4.2×10^-5^ for ACSF versus Tio at 100 pulses. **e**, Normalized representative trace of rACh1h in response to single pulse electrical stimulation. **f**, Group summary of on and off kinetics of rACh1h. mean ± s.e.m. n = 6/3 and 6/3 for on and off. **g**, Schematic illustration depicting the fiber-photometry recording involving red ACh sensors for panel **h-o**. **h**, Representative traces of rACh1h and rAChmut in response to optical stimulation in the BF before (Control, left), after an i.p. injection of the AChE inhibitor donepezil (Done, 10 mg/kg, middle), and after an i.p. injection of the M3R antagonist scopolamine (Scop, 10 mg/kg, right). **i**, Representative trace of fluorescence change in rACh1h to Done application. **j**, Group summary of fluorescence change in rACh1h to Done application. mean ± s.e.m. n = 3 mice for each rACh1h and rAChmut group. Two-tailed Student’s t tests: P = 0.005 of rACh1h before and after done application. **k**, Representative trace of fluorescence change and kinetics in rACh1h to optogenetic stimulation. **l**, Group summary of fluorescence change and decay_t50_ in rACh1h to optogenetic stimulation. mean ± s.e.m. Two-tailed Student’s t tests were performed. For peak ΔF/F_0_, P = 0.04 in rACh1h between control and Done, P = 0.0065 in rACh1h between Done and Scop. For decay_t50_, P = 6.6×10^-4^ before and after Done application. **m**, Representative traces of rACh1h and rAChmut fluorescence before and after Scop application. **n**, Normalized power spectra of the photometry signal for rACh1h and rAChmut. **o**, Left, isolated power spectrum of rACh1h and rAChmut. Right, area under the curve (AUC) for ACh power in the band at 0.02 - 1 Hz in rACh1h and rAChmut. mean ± s.e.m. Two-tailed Student’s t tests were performed, P = 0.02 for rACh1h versus rAChmut. NS, not significant.

### Using red fluorescent ACh sensors to measure ACh release *in vivo*

Next, we examined whether our red fluorescent sensors can be used to monitor ACh release *in vivo*. Previous studies found that the basolateral amygdala (BLA) receives cholinergic input from the basal forebrain (BF)^4,28^. Therefore, to determine whether rACh1h can report ACh release in the BLA *in vivo*, we injected an AAV expressing either rACh1h or rAChmut (as a negative control) into the BLA and expressed the optogenetic tool channelrhodopsin-2 (ChR2)^29^ in the BF (Fig. 2g). We then optically stimulated neurons in the BF and measured ACh signal in the BLA using fiber photometry. We found that rACh1h reliably detected both tonic ACh release and time-locked transient increases in ACh levels, with no detectable response in mice expressing rAChmut (Fig. 2h). Moreover, an i.p. injection of the acetylcholinesterase inhibitor donepezil^30^ increased both the magnitude and duration of the rACh1h signal, while an i.p. injection of scopolamine inhibited the rACh1h response (Fig. 2h-m).Notably, rACh1h’s high affinity for ACh enabled it to detect spontaneous fluctuations in ACh (Fig. 2n). Fast Fourier Transform (FFT) analysis revealed that rACh1h can report spontaneous ACh release events occurring at a frequency of 0.02-1 Hz, whereas no fluctuations were detected using rAChmut (Fig. 2o-p). We repeated these experiments using the medium-affinity and low-infinity rACh1m and rACh1l sensors (Extended Data Fig. 4a-b) and found that both sensors can reliably report the release of ACh from optically stimulated BF neurons (Extended Data Fig. 4c-h). Thus, all three red fluorescent ACh sensors can be used to monitor ACh release *in vivo* with high sensitivity and temporal resolution.

### Dual-color imaging of both ACh release and calcium signaling

We next determined whether the red fluorescent rACh1h sensor can be used together with the green fluorescent GCaMP6s sensor to simultaneously measure ACh release and changes in intracellular calcium, respectively. The modulation of medium spiny neurons (MSNs) by striatal cholinergic interneurons is critical for reinforcement learning and locomotion^31,32^. We therefore injected an AAV expressing rACh1h in the NAc while also expressing GCaMP6s in dopamine 1 receptor (D1R)–positive MSNs (Extended Data Fig. 5a). Using fiber photometry, we then recorded the rACh1h and GCaMP6s signals produced during both foot shock and reward paradigms. The results revealed that foot shock induced a robust increase in both rACh1h and GCaMP6s fluorescence, while reward induced a robust decrease in both rACh1h and GCaMP6s fluorescence, with a high correlation between the ACh and calcium signals (Extended Data Fig. 5b-e).

### Dual-color imaging of both ACh and dopamine release

Leveraging the spectral compatibility of the red ACh sensor with green fluorescent sensors, we simultaneously monitored multiple signaling molecules within the same brain region. External reward and sensory cues trigger the release of both dopamine (DA) and ACh, both of which play an important role in facilitating learning and motivation^33^. Moreover, the BLA plays a key role in associating cues with both positive and negative valence outcomes^34^. To measure the release of these two neural modulators simultaneously in the BLA, we expressed rACh1h and gDA3h in the BLA and then used fiber photometry to record ACh and DA activity, respectively, during auditory Pavlovian conditioning tasks (Fig. 3a-b). We found that rACh1h responded to both reward and punishment, whereas gDA3h responded predominantly to reward. After five days of training, both sensors exhibited a stronger response to the tone predicting reward (Fig. 3c-d). Moreover, a cross-correlation analysis revealed a high correlation between the ACh and DA signals (Fig. 3e). In addition, both the rACh1h and gDA3h signals increased in response to the conditioned stimulus following training (Fig. 3f). This development of an excitatory response to reward-predicting cues is consistent with the so-called reward-prediction-error theory^35^. Together, these results confirm that the red fluorescent rACh1h sensor is compatible for use with gDA3h, providing the ability to simultaneously measure ACh and DA release in real time.

**Fig. 3.**
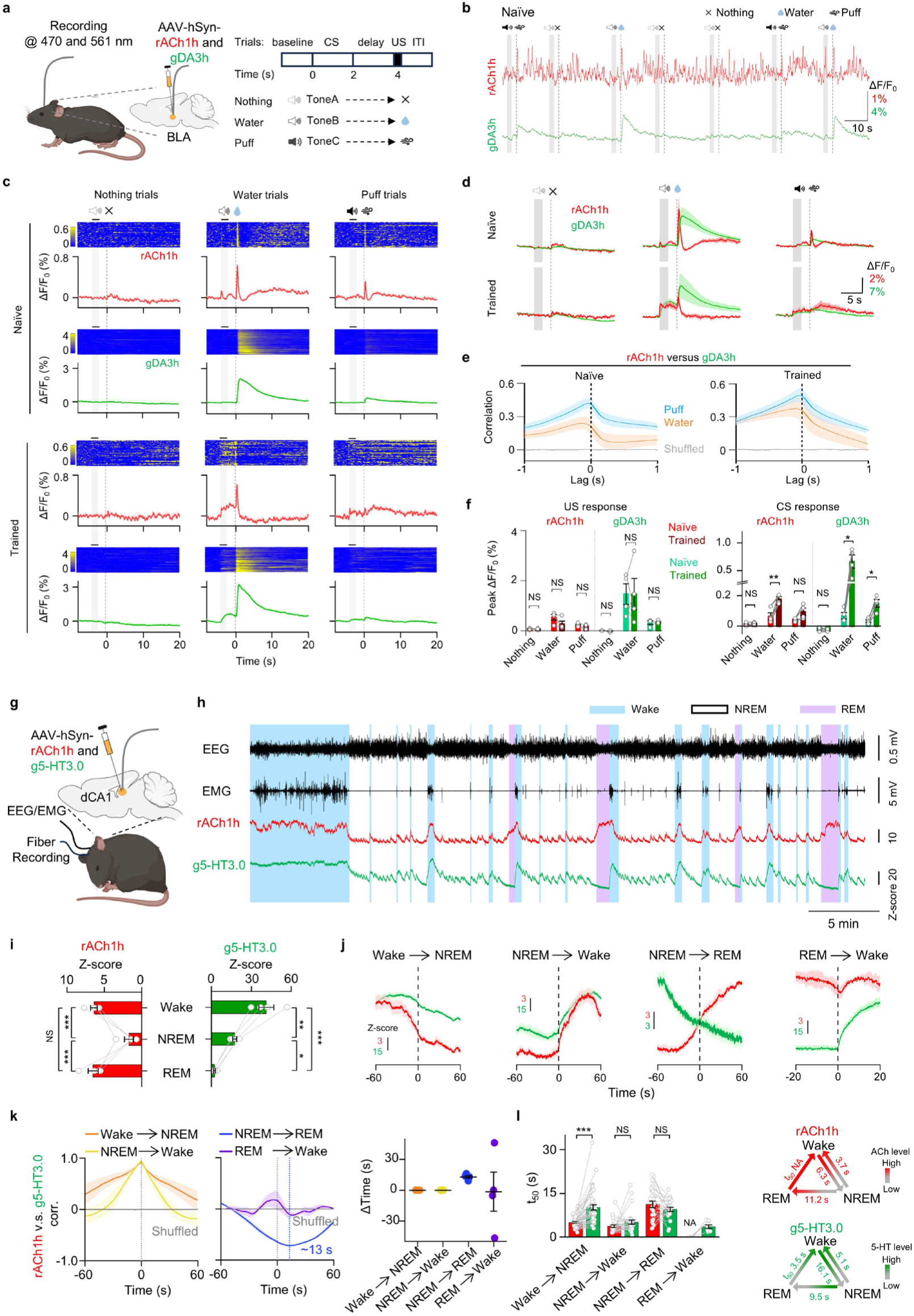
Multiplex measurements of ACh with other neuromodulators. **a**, Schematic illustration depicting the dual-color recording involving rACh1h and gDA3h during Pavlovian conditioning tasks for panel **b-f**. **b**, Representative traces of rACh1h (red) and gDA3h (green) simultaneously measured in BLA during seven consecutive trials. **c**, Representative pseudocolored images and averaged traces of rACh1h and gDA3h fluorescence from a mouse in naive (Top) and trained (Bottom) state. The gray shading indicates the application of audio. The dashed line indicates the delivery of water, puff or nothing. **d**, Group-averaged traces of rACh1h and gDA3h in the BLA for all mice under naive and trained states. mean ± s.e.m. n = 4 mice. **e**, The average cross-correlation between rACh1h and gDA3h signals under naive and trained states. **f**, Group summary of fluorescence change of rACh1h and gDA3h signals to US (left) and CS (right). mean ± s.e.m. Paired Two-tailed Student’s t tests was performed. For rACh1h, P = 0.007 in water trial. For gDA3h, P = 0.021 in water trial, P = 0.014 in puff trial. **g**, Schematic illustration depicting the dual-color recording involving rACh1h and g5-HT3.0 during sleep–wake cycles for panel **h-l**. **h**, Representative traces of EEG, EMG, rACh1h (red) and g5-HT3.0 (green) during sleep–wake cycles in freely behaving mice. Bule shading, wake state; Pink shading, REM sleep. **i**, Group summary of rACh1h and g5-HT3.0 fluorescence in dCA1 during the wake state, NREM sleep, and REM sleep. mean ± s.e.m. n = 4 mice. One-way ANOVA with post hoc Tukey’s test was performed. For rACh1h, post hoc test: P = 7.5×10^-4^ between Wake and NREM, P = 6.4×10^-4^ between NREM and REM. For g5-HT3.0, post hoc test: P = 2.2×10^-3^ between Wake and NREM, P = 5.8×10^-5^ between Wake and REM, P = 0.03 between NREM and REM. **j**, Representative time courses of rACh1h and g5-HT3.0 fluorescence signal during the indicated transitions between the sleep-wake states. **k**, Left, the average cross-correlation between rACh1h and g5-HT3.0 signals during sleep–wake cycles. Right, Group summary of time lag of cross-correlation peak between rACh1h and g5-HT3.0 signals during sleep–wake cycles. mean ± s.e.m. **l**, Group summary (left) and summary model (right) of the t50 values measured for each transition between the indicated sleep-wake states. mean ± s.e.m. Two-tailed Student’s t tests were performed. P = 2.7×10^-7^ between rACh1h and g5-HT3.0 during the transition from wake to NREM. CS, conditional stimulus; ITI, inter-trial interval; US, unconditional stimulus. NA, not available. NS, not significant.

### Simultaneously measuring ACh and serotonin release during the sleep-wake cycle

The hippocampus plays an essential role in memory consolidation during sleep and receives both cholinergic and serotonergic inputs^5,36^. To measure both ACh and serotonin (5-HT) levels during the sleep-wake cycle, we injected AAVs expressing red fluorescent rACh1h and the green fluorescent 5-HT sensor g5-HT3.0 in the dorsal CA1 region (dCA1) of the hippocampus. We then performed simultaneous fiber photometry, electroencephalography (EEG, to measure the sleep-wake cycle), and electromyography (EMG, to measure the animal’s activity) recordings in freely moving mice (Fig. 3g-h). We found that both the rACh1h and g5-HT3.0 signals were high during wakefulness, but were relatively low during non-rapid eye movement (NREM) sleep. Moreover, during rapid eye movement (REM) sleep, the rACh1h signal was high while the g5-HT3.0 signal was low (Fig. 3i), consistent with previous studies^17,21,37^. As a negative control, the ACh-insensitive rAChmut signal did not change during REM sleep (Extended Data Fig. 6). An analysis of the transition between various sleep-wake states revealed a strong positive correlation between the rACh1h and g5-HT3.0 signals during the wake-to-NREM and the NREM-to-wake transitions, and a negative correlation during the NREM-to-REM transition (Fig. 3i-k). We also calculated the t50 of the signals during transitions between sleep-wake states and found that the ACh signal decreased more rapidly compared to the 5-HT signal during the wake-to-NREM transition (Fig. 3l).

### Spatially resolved imaging of cortical ACh and norepinephrine release

Cholinergic neurons in the basal forebrain project extensively throughout the neocortex, regulating arousal, attention, and motivation^6^. Moreover, cortical activity is also shaped by input from noradrenergic neurons in the locus coeruleus^38,39^. To measure both cortical ACh and cortical norepinephrine (NE) release with high spatiotemporal resolution, we expressed both the red fluorescent rACh1h sensor and the green fluorescent NE2m sensor in the primary visual cortex (V1) and then performed head-fixed *in vivo* two-photon imaging (Fig. 4a-b). During recording, the mouse was placed on a treadmill and was exposed to a variety of stimuli, including water delivery induced by licking (Fig. 4c), flashes of light (Fig. 4d), auditory tones (Fig. 4e), and forced running (Fig. 4f). We found that during water licking, rACh1h fluorescence increased, while NE2m fluorescence was unchanged. Moreover, forced running significantly increased both rACh1h and NE2m fluorescence, whereas visual and auditory stimuli produced no response in either sensor (Fig. 4g-h). These results obtained with the rACh1h sensor are consistent with previous reports regarding the gACh3.0 signal measured in V1^17^. Interestingly, the start of the increase in rACh1h fluorescence occurred prior to the licking action, but after the start of forced running (Fig. 4i).

**Fig. 4.**
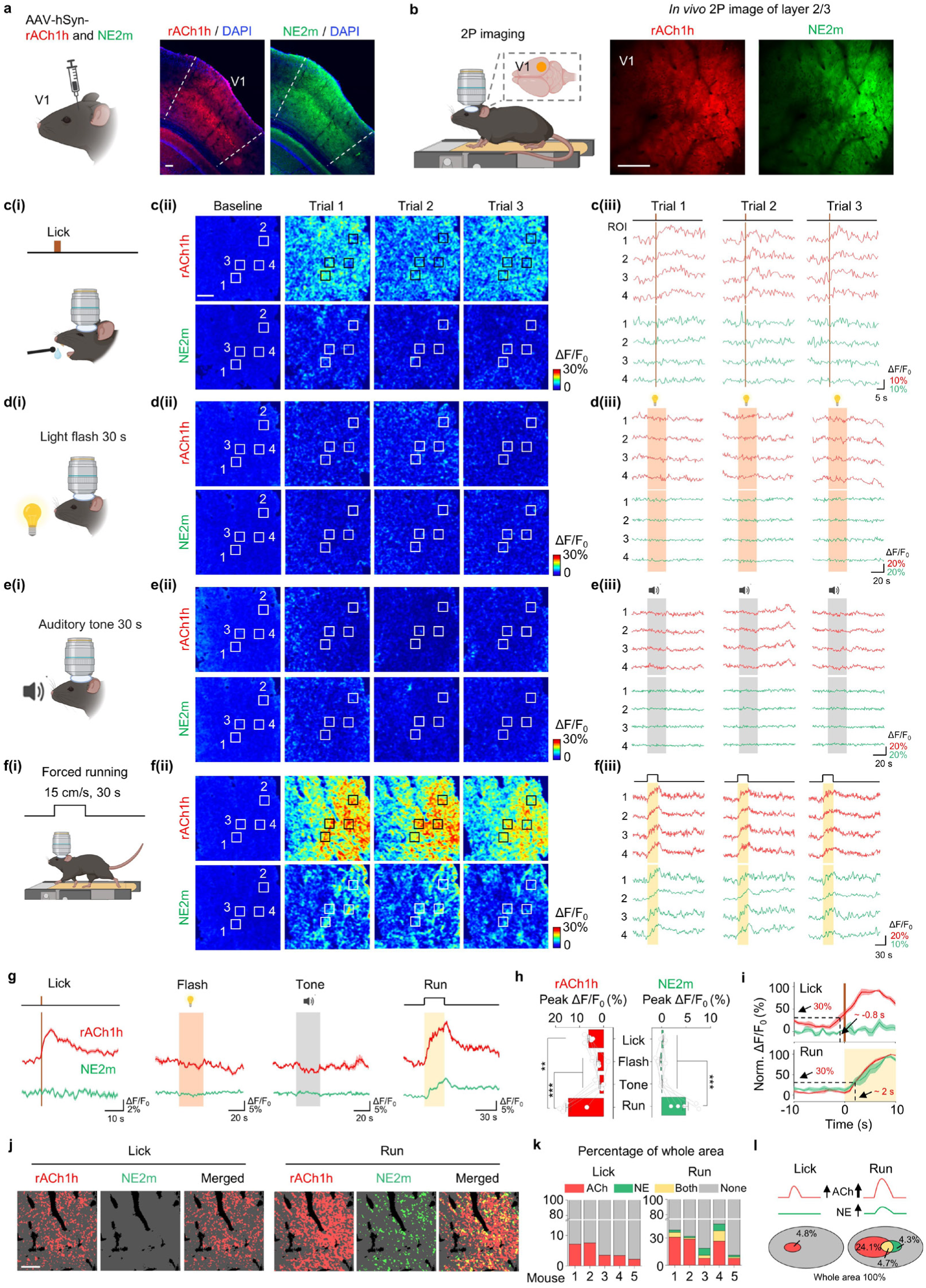
Spatially resolved cortical ACh and NE dynamics. **a**, Schematic illustration (Left and medium) depicting the viral injection and head-fixed two-photon imaging at V1 cortex. Representative image (Right) showing the expression of rACh1h and NE2m in coronal brain slice. Scale bar, 100 μm. **b**, Example in vivo two-photon images of layer 2/3 in the V1 cortex showing rACh1 and NE2m fluorescence. Scale bar, 100 μm. c, Schematic cartoon illustrating water licking task (**c(i)**), representative response images (**c(ii)**) and typical traces (**c(iii)**) during three trials for rACh1h (Top) and NE2m (Bottom). Scale bar, 100 μm. **d-f**, Similar to c, the illustration, response images and traces in light flash (**d**), auditory tone (**e**) and forced running (**f**). Two-photon imaging was performed in the same region across behaviors. **g**, Averaged traces of rACh1h and NE2m in different behaviors. **h**, Quantifications of peak ΔF/F_0_ (Left) and Z-score group summary of 5 mice (Right) for rACh1h and NE2m during different behaviors. mean ± s.e.m. n = 5 mice. One-way ANOVA with post hoc Tukey’s test was performed. For rACh1h, post hoc test: P = 1.7×10^-5^ and 8.7×10^-6^ for Run versus Tone and Flash. For NE2m, post hoc test: P = 2.4×10^-4^, 6.8×10^-5^ and 1.4×10^-4^ for Run versus Lick, Flash and Tone. **i**, Normalized trace (Left) of rACh1h and NE2m during lick and run. Arrow and dash indicating the response and time point at 30% Nrom. ΔF/F_0_. **j**, Representative images showing the special pattern for rACh1h and NE2m to lick and run. **k**, Percentage summary of the response area from 5 mice for rACh1h and NE2m to lick and run. **l**, Summary and Venn diagram for rACh1h and NE2m responding to lick and run. mean ± s.e.m. n = 5 mice. NA, not available.

Lastly, we analyzed the spatial distribution of the rACh1h and NE2m signals during water licking and forced running (Fig. 4j-l). During licking, an increase in ACh release was observed in 4.8% of the total area imaged, with no change in NE release; during running, 4.7% of the total imaged area showed an increase in both ACh and NE release, while ACh release alone and NE release alone were observed in 24.1% and 4.3%, respectively, of the total imaged area. In summary, these findings demonstrate that rACh1h can be combined with NE2m in order to simultaneously record both ACh and NE release *in vivo* with high spatiotemporal resolution.

## Discussion

Here, we developed a series of genetically encoded red fluorescent ACh sensors. We then demonstrated that these sensors can be used to monitor ACh dynamics both *in vitro* and *in vivo* with extremely high sensitivity and spatiotemporal resolution.

To maximize flexibility, we generated three versions of ACh sensors based on their ligand affinity, with rACh1h, rACh1m, and rACh1l having high, medium, and low affinity for ACh, respectively. These sensors—particularly rACh1h—exhibit a stronger response to ACh and a higher SNR compared to previously reported sensors, both in cultured cells and in cultured neurons. We then showed that rACh1h is suitable for monitoring ACh release in brain areas that receiving cholinergic input, including the basolateral amygdala, the CA1 region of the hippocampus, and V1, as well as regions containing local cholinergic neurons such as the nucleus accumbens. Importantly, the red-shifted spectrum of rACh1h allows for the simultaneous recording of ACh and a variety of other neurochemicals and signaling molecules, including calcium, dopamine, serotonin, and norepinephrine. Thus, by combining the rACh1h and g5-HT3.0 sensors, we found that ACh and 5-HT have similar oscillations during wakefulness and NREM sleep, but have opposing activity patterns during REM sleep, suggesting that these two neuromodulators have distinct roles in the brain during various sleep-wake states. Using two-photon *in vivo* imaging, we also found that rACh1h responds to both water licking and forced running—albeit with distinct times of onset—while NE2m responds only to forced running. These results suggest that ACh may regulate cortical neurons during both active and passive behaviors, whereas NE may play a more prominent role in response to passive behaviors.

The high affinity of rACh1h for ACh makes it particularly suitable for detecting spontaneous ACh release. Conversely, rACh1m and rACh1l, which have relatively lower affinity for ACh, are suitable for use in ACh-abundant brain regions such as the nucleus accumbens, producing a smaller response but with faster *τ*_off_ kinetics. This series of sensors with a range of ACh affinities greatly increase our ability to detect various changes in ACh concentration in specific brain regions.

In summary, these new red fluorescent ACh sensors significantly expand our ability to monitor ACh release with high sensitivity and spatiotemporal resolution. Moreover, their wide detection range and spectral compatibility with other fluorescent sensors provide a powerful set of tools for deciphering the complexity of the cholinergic system.

## Methods

### Animals

All animal studies and experimental procedures were approved by the laboratory animal care and use committee of Peking University. Newborn wild-type Sprague-Dawley rat pups (P0) and wild-type male C57BL/6J mice (8- to 12-weeks old, from Beijing Vital River Laboratory) were used in this study. D1R-cre mice were generously provided by Y. Rao at Peking University. All animals were group-housed or pair-housed at 18-23°C in 40-60% humidity, with a 12h/12h light/dark cycle and food and water provided ad libitum.

### AAV expression

AAV2/9-hSyn-rACh1h (9.63 × 10^13^ vg·ml^−1^), AAV2/9-hSyn-rAChmut (3.86 × 10^13^ vg·ml^−1^), AAV2/9-hSyn-rACh1m (4.40 × 10^13^ vg·ml^−1^), AAV2/9-hSyn-gACh1l (1.03 × 10^13^ vg·ml^−1^), AAV2/9-hSyn-gACh3.0 (8.0 × 10^13^ vg·ml^−1^), and AAV2/9-hSyn-iAChSnFR (3.53 × 10^13^ vg·ml^−1^) were packaged at Vigene Biosciences. AAV2/9-hSyn-hChR2(H134R)-eYFP (5.49 × 10^12^ vg·ml^−1^) and AAV2/9-DIO-hSyn-GCaMP6s (5.52 × 10^12^ vg·ml^−1^) were packaged at BrainVTA. AAV2/9-hSyn-NE2m (1.39 × 10^13^ vg·ml^−1^) was packaged at Shenzhen Bay Laboratory. Where indicated, the AAVs were either used to infect cultured neurons or injected *in vivo* into specific brain regions.

### Molecular biology

All plasmids used in this study were generated using Gibson assembly^40^, and the sequences of all clones were confirmed using Sanger sequencing. cDNAs encoding muscarinic type 3 receptors were cloned from a mouse cDNA library and a human GPCR cDNA library (hORFeome database 8.1, http://horfdb.dfci.harvard.edu/index.php?page=home). To screen and characterize the sensors in HEK293T cells, the sensor-encoding cDNAs were cloned into the pDisplay vector containing an upstream IgK leader sequence and a downstream IRES-EGFP-CAAX cassette. The EGFP-CAAX cassette provides a membrane marker and was used to calibrate fluorescence. To optimize the sensors, site-directed mutagenesis was performed using primers containing randomized NNB codons (48 codons in total, encoding all 20 possible amino acids). For expression and characterization in cultured neurons, the sensors were cloned into the pAAV vector containing the human Synapsin promoter. To measure downstream coupling using the Tango assay, the indicated rACh sensor or wild-type M_3_R was cloned into the pTango vector^26^. For the luciferase complementation assay, the β2AR gene in the β2AR-Smbit construct was replaced with the indicated rACh sensor or wild-type M_3_R; LgBit-mGs was a gift from N.A. Lambert (Augusta University).

### Cell culture

HEK293T cells were purchased from ATCC (CRL-3216). The cells were cultured at 37°C in humidified air containing 5% CO_2_ in DMEM (Biological Industries) supplemented with 10% (v/v) fetal bovine serum (Gibco) and 1% penicillin-streptomycin (Gibco). Rat cortical neurons were prepared using P0 Sprague–Dawley rat pups (both sexes) purchased from Beijing Vital River. Rat cortical neurons were dissociated from the dissected rat cerebral cortex by digestion in 0.25% trypsin-EDTA (Biological Industries) and plated on poly-D-lysine–coated (Sigma-Aldrich) 12-mm glass coverslips in 24-well plates. The neurons were cultured in Neurobasal medium (Gibco) containing 2% B-27 supplement (Gibco), 1% GlutaMAX (Gibco), and 1% penicillin-streptomycin (Gibco) at 37°C in humidified air containing 5% CO_2_.

### Fluorescence imaging of cultured cells

HEK293T cells were cultured on 12-mm glass coverslips in 24-well plates or in 96-well plates without coverslips. When the cells reached ∼70% confluence, they were transfected using PEI (1 μg plasmid and 3 μg PEI per well in 24-well plates, or 300 ng plasmid and 900 ng PEI per well in 96-well plates). Imaging was performed 24-48 h after transfection. For cortical neurons, the cells were infected with AAV expressing the indicated red fluorescent ACh sensor at 3-5 days *in vitro* (DIV3-5) and imaged at DIV11-14. Before imaging, the culture medium was replaced with Tyrode’s solution consisting of (in mM): 150 NaCl, 4 KCl, 2 MgCl_2_, 2 CaCl_2_, 10 HEPES, and 10 glucose (pH 7.4). The cells grown on coverslips were transferred to a custom-made chamber and imaged using an inverted Ti-E A1 confocal microscope (Nikon) with NIS-Element 4.51.00 software (Nikon). The confocal microscope was equipped with a 10×/0.45 numerical aperture (NA) objective, a 20×/0.75 NA objective, a 40×/1.35 NA oil-immersion objective, a 488-nm laser, and a 561-nm laser. The cells cultured in 96-well plates without coverslips were imaged using an Opera Phenix system equipped with a 20×/0.4 NA objective, a 40×/1.1 NA water-immersion objective, a 488-nm laser, and a 561-nm laser, controlled using Harmony 4.9 software.

To measure the sensors’ responses induced by various chemicals, solutions containing the following compounds were delivered to the cells by bath application or perfusion at the indicated concentrations: ACh (AMQUAR), Tio (MCE), Scop (MCE) Choline (Sigma-Aldrich), Nicotine (Tocris), glutamate (Sigma-Aldrich), GABA (γ-aminobutyric acid; Tocris), DA (Sigma-Aldrich), NE (Tocris), serotonin (Tocris), histamine (Tocris), octopamine (Tocris), tyramine (TA; Aladdin), and adenosine (Ado; Sigma-Aldrich ). The change in fluorescence (ΔF/F_0_) was measured using the formula [(F − F_0_)/F_0_], in which F_0_ is baseline fluorescence defined as the average fluorescence measured 0–1 min before drug application.

In the experiment to test whether blue light can photoactivate the red fluorescent sensors, cells expressing the indicated rACh sensors or jRGECO1a were imaged using an inverted Ti-E A1 confocal microscope, and the cells were stimulated with a 488-nm laser emitted from the objective (power: 210 μW; intensity: 0.4 W/cm^2^). The laser was applied for a duration of 1 sec.

### Spectra measurements

To measure one-photon spectra, HEK293T cells were transfected with plasmids encoding the various rACh sensors and then transferred to 384-well plates 24-30 h after transfection. The excitation and emission spectra were then measured at 5-nm increments using a Safire2 multi-mode plate reader (Tecan) in the absence and presence of 100 μM ACh. To measure two-photon spectra, HEK293T cells were plated on 12-mm coverslips, transfected with plasmid encoding the various rACh sensors, and excitation and emission spectra were measured at 10- nm increments ranging from 820-1300 nm using a FVMPE-RS microscope (Olympus) equipped with a Spectra-Physics InSight X3 dual-output laser.

### Luciferase complementation assay

HEK293T cells at 50–60% confluence were co-transfected with either wild-type M_3_R or the indicated sensor together with the corresponding LgBit-mG construct; 24–48 h post-transfection, the cells were washed in phosphate-buffered saline (PBS), dissociated using a cell scraper, and resuspended in PBS. The cells were then transferred to opaque 96-well plates containing 5 μM furimazine (NanoLuc Luciferase Assay, Promega) and bathed in ACh at concentrations ranging from 0.1 nM to 100 μM. After incubation for 10 min in the dark, luminescence was measured using a VICTOR X5 multilabel plate reader (PerkinElmer).

### Tango assay

The Tango assay was performed using the HTLA cell line, which stably expresses a tTA-dependent luciferase reporter alongside a β-arrestin2-TEV fusion gene. These cells were transfected with plasmids encoding the indicated receptors or sensors. After culturing for 24 h in 6-well plates, the cells were transferred to 96-well plates and bathed in ACh at concentrations ranging from 0.01 nM to 100 μM. After a 12-hour incubation to facilitate expression of the tTA-dependent luciferase, the Bright-Glo reagent (Fluc Luciferase Assay System, Promega) was added to a final concentration of 5 μM, and luminescence was measured using a VICTOR X5 multilabel plate reader (PerkinElmer).

### Two-photon imaging of acute mouse brain slices

Wild-type adult (6–8 weeks of age) male C57BL/6N mice were anesthetized with an i.p. injection of 2,2,2-tribromoethanol (Avertin, 500 mg per kg body weight; Sigma-Aldrich), and AAV-hSyn-rACh1h was injected (300 nl at a rate of 50 nl·min^−1^) into the NAc using the following coordinates: anterior-posterior (AP): +1.4 mm relative to Bregma; medial-lateral (ML): ±1.2 mm relative to Bregma; and dorsal-ventral (DV): −4.0 mm from the dura. Two weeks after virus injection, the mice were deeply anesthetized, and the heart was perfused with slicing buffer containing (in mM): 110 choline chloride, 2.5 KCl, 1.25 NaH_2_PO_4_, 25 NaHCO_3_, 7 MgCl_2_, 25 glucose, and 0.5 CaCl_2_. The mice were then decapitated, and the brains were immediately removed and placed in cold oxygenated slicing buffer. The brains were sectioned into 300-μm-thick coronal slices using a VT1200 vibratome (Leica), and the slices were incubated at 34°C for at least 40 min in oxygenated artificial cerebrospinal fluid containing (in mM): 125 NaCl, 2.5 KCl, 1 NaH_2_PO_4_, 25 NaHCO_3_, 1.3 MgCl_2_, 25 glucose, and 2 CaCl_2_. Two-photon imaging was performed using an Ultima Investigator two-photon microscope (Bruker) equipped with a 20×/1.00 NA objective (Olympus) and an InSight X3 tunable laser (Spectra-Physics), using Prairie View 5.5 software (Bruker). A 1040-nm laser was used to excite the rACh1h sensor, and a 595/50-nm emission filter was used to collect the fluorescence signal. For electrical stimulation, a bipolar electrode (model WE30031.0A3, MicroProbes) was positioned near the NAc core under fluorescence guidance, and imaging and stimulation were synchronized using an Arduino board with custom-written software; the stimulation voltage was set at 4–5 V. Where indicated, compounds were added by perfusion at a flow rate of 4 ml·min^−1^.

### Fiber photometry recording of optogenetically induced ACh release in mice

Adult male C57BL/6J mice, aged 8-9 weeks, were used in this study. They were anesthetized using 1.5% isoflurane and secured in a stereotaxic apparatus to ensure precise targeting during the procedure. AAV-hSyn-rACh1h, AAV-hSyn-rACh1m, AAV-hSyn-rACh1l or AAV-hSyn-rAChmut (300 nl) was injected into the BLA using the following coordinates: AP: -1.4 mm relative to Bregma; ML: ±3.0 mm relative to Bregma; and DV: -4.0 mm from the dura. For activation of the BF, AAV-hSyn-ChR2-YFP (300 nl) was injected into the BF using the following coordinates: AP: 0 mm relative to Bregma; ML: ±1.5 mm relative to Bregma; and DV: -4.8 mm from the dura. Two optical fibers (200-μm diameter, 0.37 NA; Inper) were implanted; one optical fiber was positioned 0.1 mm above the virus injection site in the BLA to record the ACh sensor, while the other optical fiber was positioned 0.3 mm above the virus injection site in the BF to optically activate ChR2. The optical fibers were secured to the skull surface using dental cement (3M).

Two to three weeks after vector injection, fluorescence signals were recorded using a fiber photometry system (FPS-410/470/561; Inper). Yellow light-emitting diode (LED) light was bandpass-filtered (561/10 nm), reflected by a dichroic mirror (495 nm), and then focused using a 20× objective (Olympus). An optical fiber was used to guide the light between the commutator and the implanted optical fiber cannulas. The excitation light emitted by the LED was set to 20-30 μW and delivered at 10 Hz with a 10-ms pulse duration. The optical signals were then collected through the optical fibers. Red fluorescence was bandpass-filtered (520/20 nm and 595/30 nm) and captured using an sCMOS camera. The current output generated by the photomultiplier tube was transduced into a voltage signal using an amplifier (A-M Systems) and subsequently passed through a low-pass filter to remove high-frequency noise. The analog voltage signals were then digitized using an acquisition card (National Instruments). To reduce autofluorescence generated by the optical fibers, the recording fibers were photobleached using a high-power LED before recording. Background autofluorescence was recorded and subtracted from the recorded signals in the subsequent analysis. A 488-nm laser (1-160 mW, LL-laser) was used for optical stimulation, with the light power at the fiber tip set at 10 mW. Optical stimuli were delivered at 40 pulses with 10ms duration for 1 s concurrently with photometry recording. Where indicated, the mice received an i.p. injection of donepezil (3 mg per kg body weight) followed by an i.p. injection of scopolamine (10 mg per kg body weight).

The photometry data were analyzed using a custom-written MATLAB program (MATLAB R2022a, MathWorks). To calculate ΔF/F_0_, baseline fluorescence (F_0_) was defined as the average fluorescence measured 5 s before the five trials of optical stimulation under control conditions.

### Dual-color recording of Calcium and rACh1h in the NAc

Adult male and female D1R-Cre mice (10-14 weeks old) were used for this experiment. AAV9-hsyn-rACh1h and AAV9-hsyn-DIO-GCaMP6s (1:1 mix, 500 nl total volume) was unilaterally injected into the NAc (AP: +1.4 mm relative to Bregma, ML: ±1.2 mm relative to Bregma, and DV: -4.0 mm from the dura), and an optical fiber (200-μm diameter, 0.37 NA; Inper) was implanted 0.1 mm above the virus injection site. Photometry recording was performed 2-3 weeks after virus injection using a customized three-color photometry system (Thinker Tech). A 470/10-nm (model 65144; Edmund optics) filtered LED at 40 μW was used to excite the green fluorescent sensors; and a 555/20-nm (model ET555/20x; Chroma) filtered LED at 40 μ W was used to excite the red fluorescent sensors; The excitation lights were delivered sequentially at 20-Hz with a 10-ms pulse duration for each, and fluorescence was collected using an sCMOS (Tucsen) and filtered with a three-bandpass filter (model ZET405/470/555/640m; Chroma). To minimize autofluorescence from the optical fiber, the recording fiber was photobleached using a high-power LED before recording.

An intraoral cheek fistula was implanted in each mouse for sucrose delivery. Incisions were made in the cheek and the scalp at the back of the neck. A short, soft silastic tube (inner diameter: 0.3 mm; outer diameter: 0.7 mm) connected via an L-shaped stainless-steel tube was then inserted into the cheek incision site. The steel tube was routed through the scalp incision, with the opposite end inserted into the oral cavity. After 3 d of recovery from the surgery, the mice were water-restricted for 36 h (until reaching 85% of their initial body weight). The water-restricted, freely moving mice then received 5% sucrose water delivery (approximately 8 μl per trial, with 25-50 trials per session and a trial interval of 20-30 s).

Before foot shock, the mice were placed in a shock box and habituated for 30 min. During the experiment, 10 1-s pulses of electricity were delivered at 0.7 mA, with an interval of 90-120 s between trials.

### Fiber photometry recordings and polysomnographic recordings during the sleep-wake cycle

Adult wild-type C57BL/6J mice were anesthetized with isoflurane and placed on a stereotaxic frame for AAV injection (400 nl per site). For the experiments shown in Fig. 3a-f, a combination of AAV-hSyn-rACh1h and AAV-hSyn-g5-HT3.0 was injected into the dCA1 using the following coordinates: AP: -2.0 mm relative to Bregma; ML: ±1.5 mm relative to Bregma; and DV: -1.4 mm from the dura. For the experiments shown in Extended Data Fig. 5, a combination of AAV-hSyn-rAChmut and AAV-hSyn-g5-HT3.0 was injected into the dCA1 using the coordinates indicated above. An optical fiber cannula (200-μm diameter, 0.37 NA; Inper) was placed 0.1 mm above the virus injection site to record the sensor signals and was affixed to the skull using dental cement.

To monitor the animal’s sleep-wake state, custom-made EEG and EMG electrodes were attached and affixed to the skull via a microconnector. The EEG electrodes were implanted into craniotomy holes positioned above the frontal cortex and visual cortex, while the EMG wires were placed bilaterally in the neck musculature. The microconnector was attached securely to the skull using glue and a thick layer of dental cement. After surgery, the mice were allowed to recover for at least 2 weeks.

The same fiber photometry system (Inper) was used to record the fluorescence signals in freely moving mice during the sleep-wake cycle. For the experiments shown in Fig.3g-h and Extended Data Fig.6a-c, a 10-Hz 470/10-nm filtered light (20-30 μW) was used to excite the green fluorescent 5-HT sensor, and a 561/10-nm filtered light (20-30 μW) was used to excite the red fluorescent ACh sensors. The fluorescent signals were captured using a dual-band bandpass filter (520/20 nm and 595/30 nm), with 10-ms pulses of excitation light delivered at 10 Hz.

The photometry data were analyzed using a custom MATLAB program. To calculate ΔF/F_0_ during the sleep-wake cycle, baseline values of the ACh signal were measured during NREM sleep, while baseline values of the 5-HT signal were measured during REM sleep. To compare the change in fluorescence between animals, ΔF/F_0_ was divided by the standard deviation of the baseline signal in order to obtain a *z*-score.

### Polysomnographic recording and analysis

The animal’s sleep-wake state was determined using EEG and EMG recordings. The signals were amplified (NL104A, Digitimer), filtered (NL125/6, Digitimer) at 0.5-100 Hz (for EEG) or 30-500 Hz (for EMG), and then digitized using a Power1401 digitizer (Cambridge Electronic Design Ltd.). Recordings were performed using Spike2 software (Cambridge Electronic Design Ltd.) at a sampling rate of 1000 Hz. The sleep-wake state was classified semi-automatically in 4-s epochs using AccuSleep and then manually confirmed using a custom-made MATLAB GUI. The wake state was defined as desynchronized low-amplitude EEG activity and high-amplitude EMG activity. NREM sleep was defined as synchronized EEG activity with high-amplitude delta frequencies (0.5-4 Hz) and low EMG activity. REM sleep was defined as prominent theta frequencies (6-10 Hz) combined with low EMG activity. EEG spectral analysis was performed using a short-time Fast Fourier Transform (FFT).

### Pavlovian auditory conditioning task

Adult (8-9 weeks of age) male C57BL/6J mice were used for these experiments. A mixture of AAV-hSyn-rACh1h (200 nl) and AAV-hysn-gDA3h (200 nl) was injected into the right BLA as described above. An optical fiber cannula (Inper) was then implanted 0.1 mm above the virus injection site in the BLA to record the ACh and DA signals. A stainless-steel head holder was attached to the skull using resin cement to head-fix the animal. An intraoral cheek fistula was then implanted in each mouse for water delivery as described above. Head-fixed mice were habituated to the treadmill apparatus for 2 d (1 h per day) prior to the experiments in order to minimize stress. The mice were water-restricted for 36 h (until reaching 85% of their initial body weight). On the day of the experiment, the Pavlovian auditory conditioning task was performed using three pairs of auditory cues and outcomes: tone A (2.5 kHz, 70 dB, 2-s duration) was paired with delivery of 10 μl of 5% sucrose water; tone B (15 kHz, 70 dB, 2-s duration) was paired with delivery of an air puff to the eye; and tone C (7.5 kHz, 70 dB, 2-s duration) was paired with no delivery. These three pairs were randomly delivered for a total of 300 trials, with a 20-30-s inter-trial interval. The delivery of water and air puff was precisely controlled by a stepper motor pump and solenoid valve, respectively. A custom-written Arduino code was used to control the timing of the pump and solenoid valve, and to synchronize the training devices with the photometry recording system.

Two weeks after virus injection, the same fiber photometry system (Inper) was used to capture the fluorescence signals. In brief, a 10-Hz (10-ms pulse duration) 470/10-nm filtered LED at 20-30 μW was used to excite the green fluorescent sensors, and a 10-Hz (10-ms pulse duration) 561/10-nm filtered LED at 20-30 μW was used to excite the red fluorescent sensors. Alternating excitation wavelengths were delivered, and fluorescence signals were collected using a sCMOS camera during dual-color imaging. To calculate ΔF/F_0_, baseline fluorescence (F_0_) was defined as the average fluorescence signal measured 4.5-5.0 s before the first auditory cue.

### Two-photon *in vivo* imaging in mice

Adult (6–8 weeks of age) male C57BL/6N mice were anesthetized with an i.p. injection of 2,2,2-tribromoethanol (Avertin, 500 mg per kg body weight; Sigma-Aldrich), the scalp was retracted, and the skull above the primary visual cortex (V1) was removed. A mixture of AAVs expressing rACh1h and NE2m (1:1 mixture, 300 nl total volume, full titer) was injected into V1 using the following coordinates: AP: -2.5 mm relative to Bregma; ML: 2.2 mm relative to Bregma; and DV: -0.5 mm from the dura. A 3.0-mm diameter round coverslip was used to replace the missing skull section. A stainless-steel head holder was attached to the skull to head-fix the animal and to reduce motion-induced artifacts during imaging. Three weeks after virus injection, wake mice were habituated for 15 min in the treadmill-adapted imaging apparatus in order to minimize stress. The motor cortex was imaged at a depth of 100–200 μm below the pial surface using Prairie View 5.5.64.100 software with an Ultima Investigator two-photon microscope (Bruker) equipped with a 16×/0.80 NA water-immersion objective (Olympus) and an InSight X3 tunable laser (Spectra-Physics). An interlaced scan pattern model with a 920-nm tunable laser and a 1040-nm fixed laser was used for sequential excitation. A 525/70-nm emission filter for NE2m and a 595/50-nm emission filter for rACh1h were used to collect the fluorescence signal. For the water licking paradigm, the animals were water-restricted for 2 days before imaging. During 2P in vivo imaging, water was provided to the mouse 1 s after the initial lick and withheld for the subsequent 60 s, regardless of further licking. A custom Arduino code was used to record the capacitance changes due to licking and to control the water delivery. For flash stimulation, 30 pulses (0.2-s duration and 0.8-s interval) of white light were delivered. For auditory stimulation, a 30-s 7000-Hz tone at 80 dB was delivered. For forced running, running speed was set at 15 cm·s^−1^. For image analysis, motion-related artifacts were corrected using EZcalcium^41^. Fluorescence intensity was measured at the indicated regions of interest (ROIs) using ImageJ software. To measure ΔF/F_0_, F_0_ was defined as the average baseline fluorescence signal measured for 10 s before the behavior onset. A *z*-score was calculated dividing ΔF/F_0_ by the standard deviation of the baseline. The peak response during a behavior was calculated as the average signal measured for 10 s (or 3 s for licking) around the maximum ΔF/F_0_ achieved after the behavior onset. For the area analysis in Fig. 4j-k, a given brain area was deemed to be responsive if the average SNR in a 10-s window (for running) or a 3-s window (for licking) surrounding the peak exceeded 1.5x the value.

### Immunohistochemistry

Mice were anesthetized with Avertin and intracardially perfused with PBS followed by 4% paraformaldehyde (PFA) in PBS, and the brains were dissected and fixed at 4°C overnight in 4% PFA in PBS. The brains were then sectioned at 40-μm thickness using a VT1200 vibratome (Leica). The slices were placed in blocking solution containing 5% (v/v) normal goat serum, 0.1% Triton X-100, and 2 mM MgCl_2_ in PBS for 30 min at room temperature. The slices were then incubated overnight at 4°C in blocking solution containing 0.5% (v/v) normal goat serum, 0.1% Triton X-100, and 2 mM MgCl_2_ in PBS with anti-GFP antibody (Abcam, catalog no. ab13970, chicken, dilution 1:500) and anti-mCherry antibody (Abcam, catalog no. ab125096, mouse, dilution 1:500). The following day, the slices were rinsed three times in blocking solution and then incubated for 2 h at room temperature with the following secondary antibodies: goat anti-chicken Alexa Fluor 488 (Abcam, catalog no. ab150169, dilution 1:1000) and goat anti-rabbit iFluor 555 (AAT Bioquest, catalog no. 16690, dilution 1:1000). After three washes in PBS, the slices were incubated in PBS containing DAPI (MedChemExpress, catalog no. HY-D0814, 5 mg·ml^−1^, dilution 1:1,000) for 15 min at room temperature, rinsed in PBS, mounted on slides, and imaged using an Aperio VERSA slide scanner (Leica) equipped with a 10× objective.

### Statistics

Except where indicated otherwise, all summary data are presented as the mean ± s.e.m. Imaging data were analyzed using ImageJ version 1.53c or MATLAB R2020a and R2022a. Group data were plotted using OriginPro 2020b (OriginLab), or Prism 8.0.2 (GraphPad). The SNR was calculated by dividing the peak response by the standard deviation of the baseline fluorescence. Differences were analyzed using the two-tailed Student’s *t*-test or one-way ANOVA; \**P* < 0.05, \*\**P* < 0.01, \*\*\**P* < 0.001, and NS, not significant (*P* ≥ 0.05).

## Acknowledgements

This work was supported by grants from the Space Medical Experiment Project of CMSP (grant nos. HYZHXMN01009), the National Natural Science Foundation of China (31925017), the National Key R&D Program of China (2022YFE0108700 and 2023YFE0207100), Beijing Municipal Science & Technology Commission (Z220009), and the NIH BRAIN Initiative (1U01NS120824) to Y.L., and National Natural Science Foundation of China (82271210, 82471214) to C.W. Support was also provided by the Feng Foundation of Biomedical Research, the New Cornerstone Science Foundation through the New Cornerstone Investigator Program, the Peking-Tsinghua Center for Life Sciences, the State Key Laboratory of Membrane Biology at Peking University School of Life Sciences (to Y.L.).

We thank Xiaoguang Lei at PKU-CLS, the optical Imaging platform, and small animal imaging platform of National Center for Protein Sciences at Peking University in Beijing, China, for their support and assistance with the Opera Phenix, the Operetta CLS high-content imaging system, the Nikon A1RSi+ laser scanning microscope, and the behavior facility. Cartoon illustrations, including Figs. 2a, 2g, 3a, 3g, 4a, 4b, 4c(i), 4d(i), 4e(i), 4f(i), Extended Data Figs. 4a, 5a, 6a, 7a(i), and 7b(i), were created with BioRender.com.

## Author contributions

Y.L. supervised the study. S.X., J.F., and Y.L. designed the study. G.L., M.L., R.C., and L.G. performed the experiments related to the development, optimization and characterization of the sensors in cultured HEK293T cells and in neurons. S.X. and E.J. performed the surgery and two-photon imaging experiments related to the validation of the sensors in acute brain slices. X.M. and J.W. performed the in vivo fiber photometry recoding during optogenetic stimulation under the supervision of C.W.. Y.Z. performed the in vivo fiber photometry recording in the NAc during foot shock and sucrose water delivery. X.M. performed the in vivo fiber photometry recording in the BLA during Pavlovian conditioning task and in the dCA1 during sleep-wake cycle. S.X. performed the in vivo two-photon imaging of the V1 cortex. S.L. performed immunohistochemistry experiments. All authors contributed to the interpretation and analysis of the data. S.X. and Y.L. wrote the manuscript with contributions from all authors.

---

**Extended Data Fig.1.**
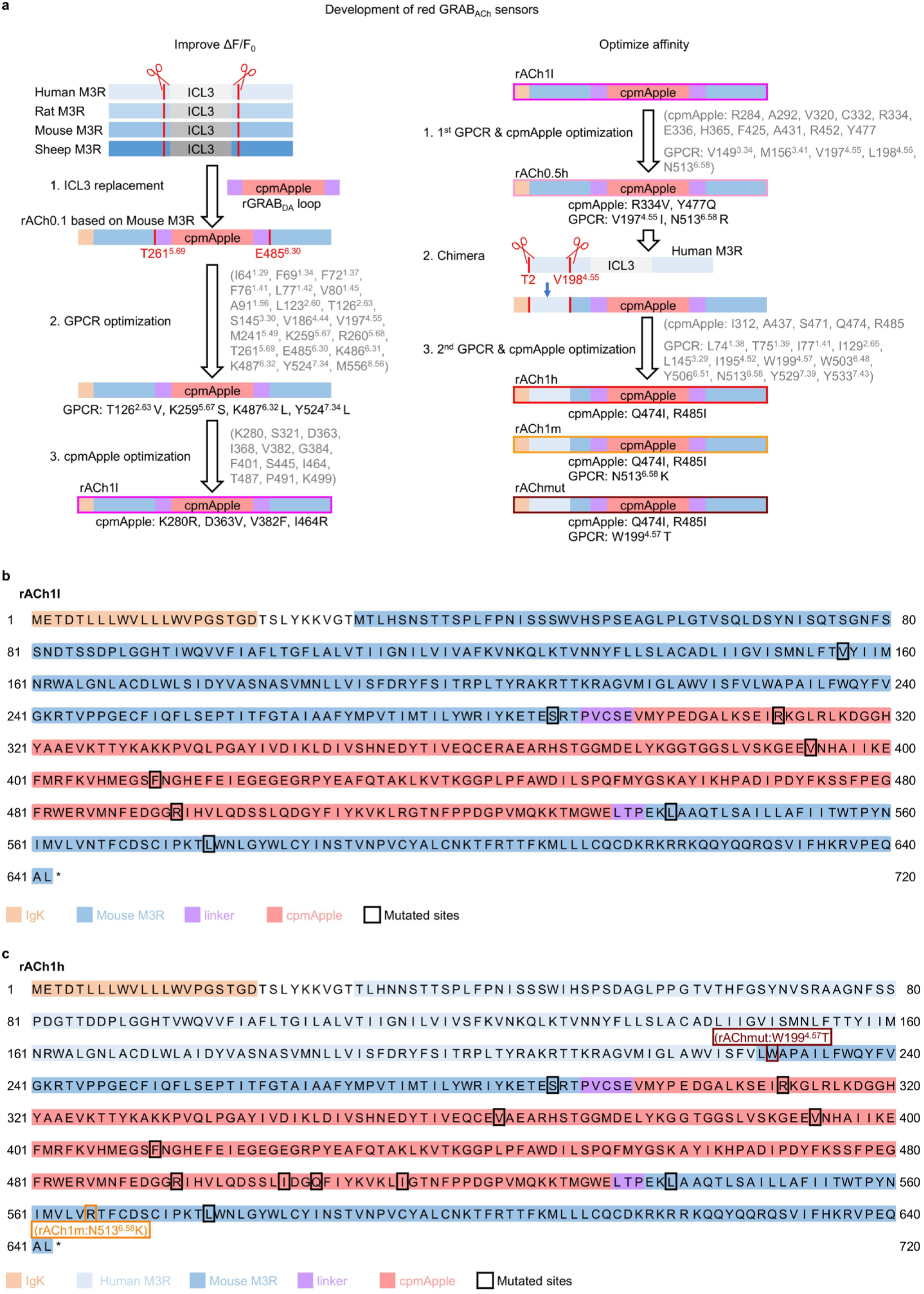
Strategy for developing and screening the red GRAB_ACh_ sensors. **a**, A flowchart showing the improving ΔF/F_0_ (Left) and optimizing affinity (Right) in the development process of the red GRAB_ACh_ sensors. The ICL3 domain of Mouse M3R was replaced by the entire ICL3 (including linker and cpmApple) derived from rGRAB_DA_. Newly generated candidate with highest ΔF/F_0_ after ICL3 replacement was then selected for further GPCR optimization and cpmApple engineering. The amino acids in gray indicating the sites screened and black indicating the sites fixed for the final candidate. In the optimization of affinity, the fragment (from T2 to V198) was replaced into rACh0.5h candidate for chimeric red ACh sensors. The candidate with highest ΔF/F_0_ was then screened for tunning the affinity to obtain rACh1h and rACh1m. **b-c**, Amino acids sequence of rACh1l (Top) and rACh1h (Bottom). The mutations adopted in the red sensors are indicated by the black box. The arginine residue at position 513^6.58^ in the mouse M3R was mutated to lysine to generate the rACh1m sensor (indicated by the orange box). The tryptophan residue at position 199^4.57^ in the mouse M3R was mutated to threonine to generate the rAChmut sensor (indicated by the dark red box).

**Extended Data Fig.2.**
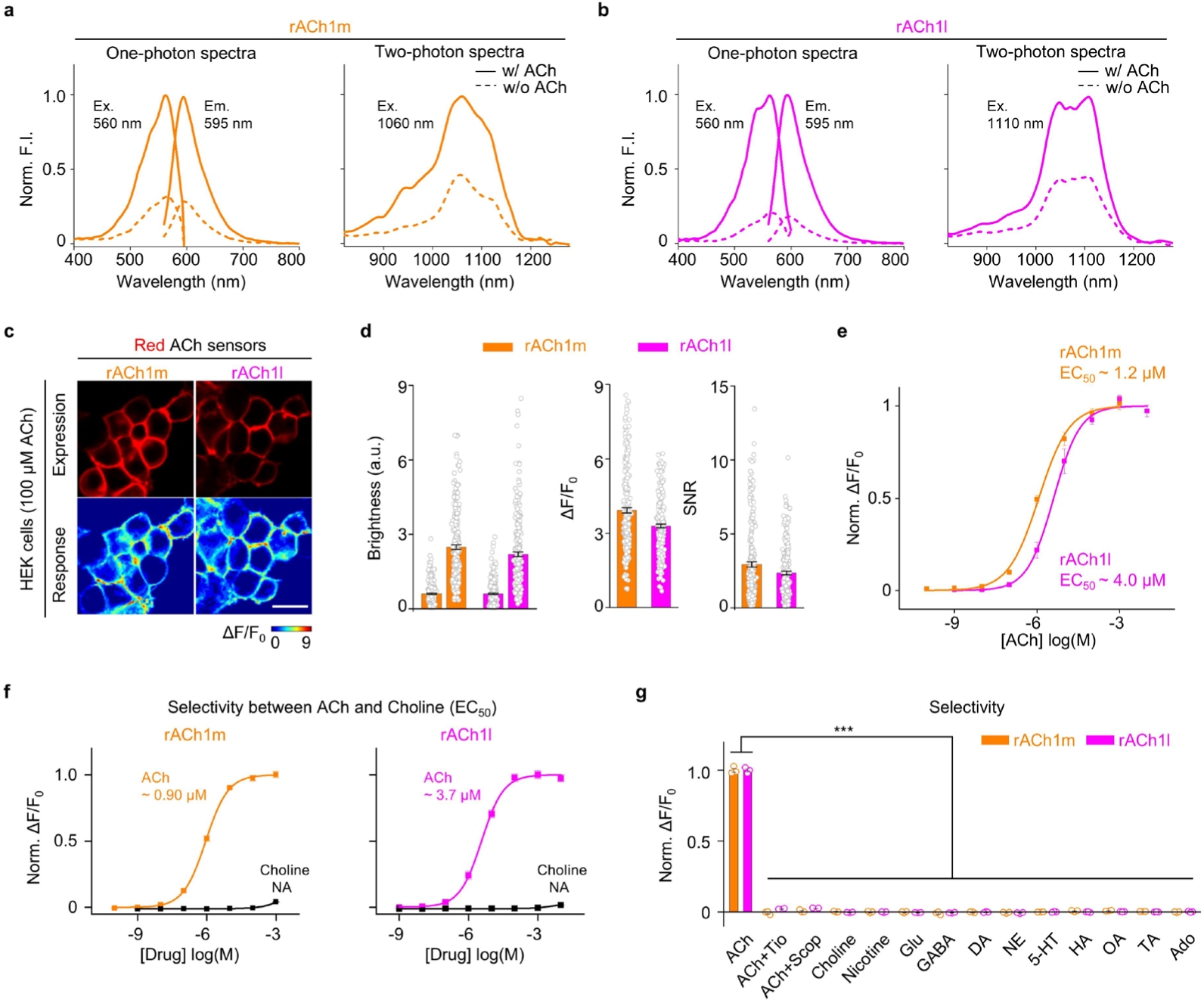
Performance of GRAB_ACh_ sensors in HEK293T cells. **a-b**, One photon and two photon spectral profiles of rACh1m (**a**) and rACh1l (**b**) in the absence (dashed line) or presence of 10 μM ACh (solid line). Left, one photon excitation and emission spectra. Right, two photon spectra of green and red ACh sensors. **c**, Representative images showing expression (top) and response (bottom) to 100 μM ACh of rACh1m and rACh1l. Scale bar, 100 μm. **d**, quantification of brightness, ΔF/F_0_ and signal-to-noise ratio of rACh1m and rACh1l before and after 100 μM ACh addition, mean ± s.e.m. n = 240/6 for each sensor. **e**, Normalized dose-response curves of rACh1m and rACh1l. n = 3 wells for each sensor. **f**, Dose– response curves for ACh and choline of rACh1m and rACh1l. n = 3 wells with 300 - 500 cells per well each. **g**, Pharmacological specificity of rACh1m and rACh1l in HEK cells. Tiotropium (Tio), M3R antagonist; Scopolamine (Scop), M3R antagonist; Glu, glutamate; GABA, γ-aminobutyric acid; DA, dopamine; NE, norepinephrine; 5-HT, serotonin; HA, histamine; OA, octopamine; TA, tyramine; Ado, adenosine. Antagonists were applied at 100 μM, others at 10 μM. n = 3 wells for rACh1m and rACh1l, 300 - 500 cells per well, mean ± s.e.m. One-way ANOVA with post hoc Tukey’s test was performed. For rACh1m and rACh1l, post hoc test: P = 0 for ACh versus ACh + Tio, ACh + Scop and other compounds. **h**, Representative traces in response to blue light in cells expressing jRGECO1a, rACh1m and rACh1l. **i**, peak ΔF/F_0_ in response to blue light in cells expressing jRGECO1a, rACh1m and rACh1l. n = 82/5, 53/5 and 67/5 for jRGECO1a, rACh1m and rACh1l. Data of jRGECO1a was replotted from Fig.1g. One-way ANOVA with post hoc Tukey’s test was performed, post hoc test: P = 0 for jRGECO1a versus rACh1m and P = 0 for jRGECO1a versus rACh1l. EC_50_, half-maximum effective concentration; FI, fluorescence intensity; NS, not significant.

**Extended Data Fig.3.**
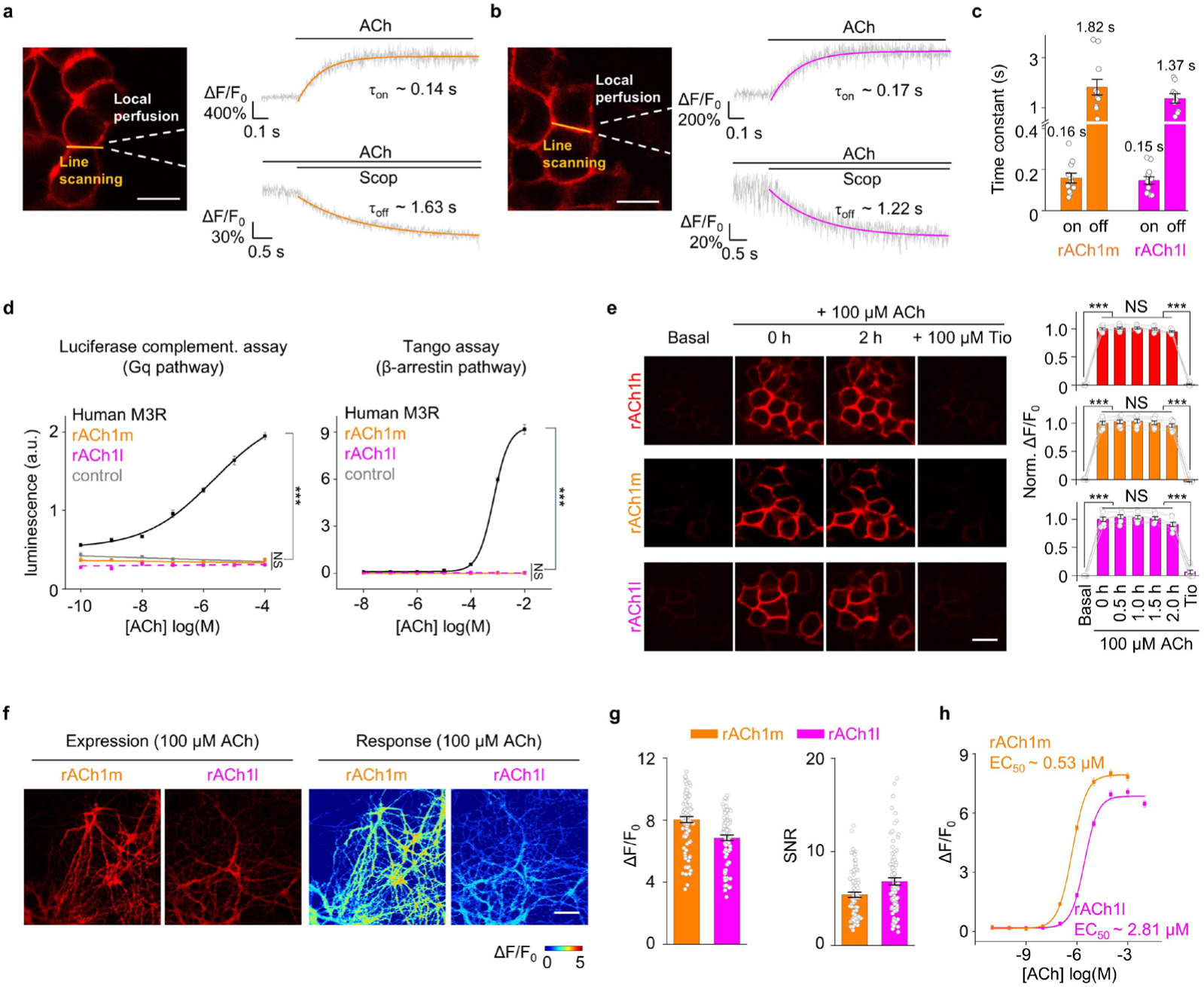
Characterization of GRAB_ACh_ sensors in cultured cells and neurons. **a-b**, Kinetics measurements of rACh1m and rACh1l. Schematic illustration showing the experimental setup of line-scanning and local puffing (Left), representative traces of sensor fluorescence increase in response to ACh (Right top) and decrease in response to Scop (Right bottom). **c**, Group summary of on and off kinetics for the sensors, mean ± s.e.m. n = 11/3 for rACh1m on kinetics; n = 11/3 for rACh1m off kinetics. n = 12/3 for rACh1l on kinetics; n = 9/3 for rACh1l off kinetics. **d**, Downstream coupling test. Human M3R; rACh1m; rACh1l; Control, without expression of WT M3R or sensors. n = 3 wells per group, One-way ANOVA with post hoc Tukey’s test was performed. For the luciferase complementation assay, post hoc test: P = 0 for rACh1m versus Human M3R, P = 0 for rACh1l versus Human M3R. For the tango assay, post hoc test: P = 0 for rACh1m versus Human M3R, P = 0 for rACh1l versus Human M3R. **e**, Representative images and normalized ΔF/F_0_ of rACh1h, rACh1m and rACh1l in response to 100 μM ACh addition, followed by 100 μM Tio. N = 7,8 and 8 well for rACh1h, rACh1m and rACh1l, mean ± s.e.m. One-way ANOVA with post hoc Tukey’s test was performed. For rACh1h, P = 0 between baseline and 0 h; P = 0 between 2 h and Tio; For rACh1m, P = 0 between baseline and 0 h; P = 0 between 2 h and Tio; for rACh1l, P = 6.5×10^-8^ between baseline and 0 h; P = 4.3×10^-8^ between 2 h and Tio. **f**, Expression and response of rACh1m and rACh1l in cultured neurons with 100 μM ACh addition. Scale bar, 100 μm. **g**, Group summary of ΔF/F_0_ and SNR of rACh1m and rACh1l. n = 90/3 for each sensor, mean ± s.e.m. **h**, Dose-response curves for rACh1m and rACh1l. n = 90/3 for each sensor. EC_50_, half-maximum effective concentration; NS, not significant.

**Extended Data Fig.4.**
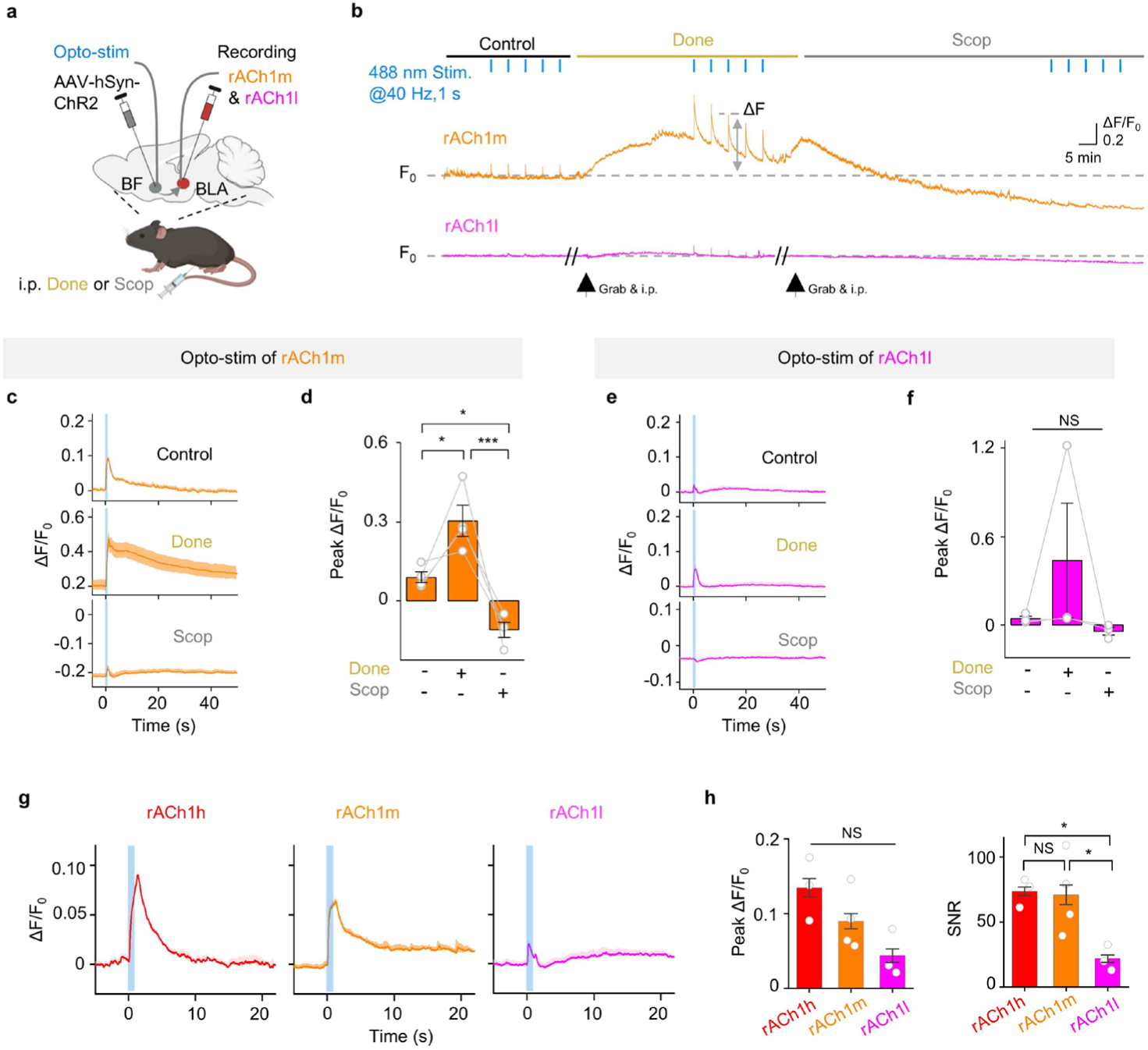
Performance of red GRAB_ACh_ sensors in vivo. **a**, Schematic illustration depicting the fiber-photometry recording involving red ACh sensors for panel **b-f**. **b**, Representative traces of rACh1m and rACh1l in response to optical stimulation in the BF before (Control, left), after an i.p. injection of the AChE inhibitor donepezil (Done, 10 mg/kg, middle), and after an i.p. injection of the M3R antagonist scopolamine (Scop, 10 mg/kg, right). **c**, Representative trace of fluorescence change of rACh1m to optogenetic stimulation. **d**, Group summary of fluorescence change in rACh1m to optogenetic stimulation. mean ± s.e.m. n = 4 mice for rACh1m and n = 3 mice for rACh1l. One-way ANOVA with post hoc Tukey’s test was performed, post hoc test: P = 0.011 for control versus Done; P = 0.016 for control versus Scop; P = 1.3×10^-6^ for Scop versus Done. **e**, Representative trace of fluorescence change of rACh1l to optogenetic stimulation. **f**, Group summary of fluorescence change in rACh1l to optogenetic stimulation. **g**, Representative trace of fluorescence change of red ACh sensors to optogenetic stimulation. Data of rACh1h is replotted from Fig.2k. **h**, Group summary of fluorescence change in red ACh sensors to optogenetic stimulation. mean ± s.e.m. n = 3 mice for rACh1h, n = 4 mice for rACh1m and n = 3 mice for rACh1l. One-way ANOVA with post hoc Tukey’s test was performed. For SNR, post hoc test: P = 0.047 for rACh1h versus rACh1l, P = 0.044 for rACh1m versus rACh1l. SNR, signal to noise ratio. NS, not significant.

**Extended Data Fig. 5.**
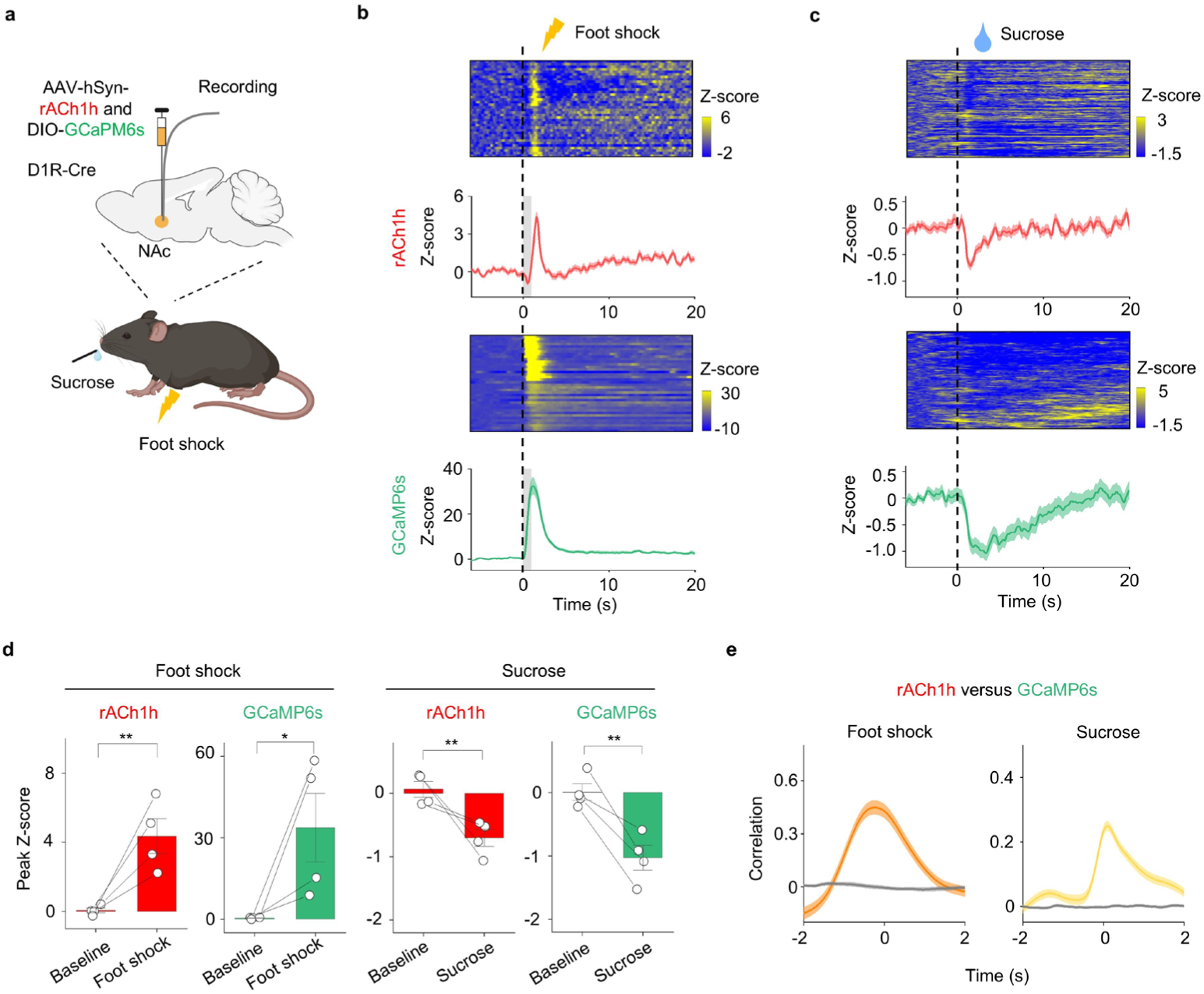
Multiplex measurements of ACh with calcium. **a**, Schematic illustration depicting the multiplex recording of rACh1h and GCaMP6s in foot shock and reward task. **b-c**, Representative pseudocolored images and averaged traces of rACh1h and GCaMP6s fluorescence from 4 mice in foot shock and reward task. **d**, Group summary of fluorescence change of rACh1h and GCaMP6s signals. n = 4 mice for foot shock and reward. mean ± s.e.m. Two-tailed Student’s t tests were performed. P = 0.006 of rACh1h and P = 0.039 of GCaMP6s before and after foot shock; P = 0.006 of rACh1h and P = 0.005 of GCaMP6s before and after reward. **e**, The average cross-correlation between rACh1h and GCaMP6s signals during foot shock and reward.

**Extended Data Fig. 6.**
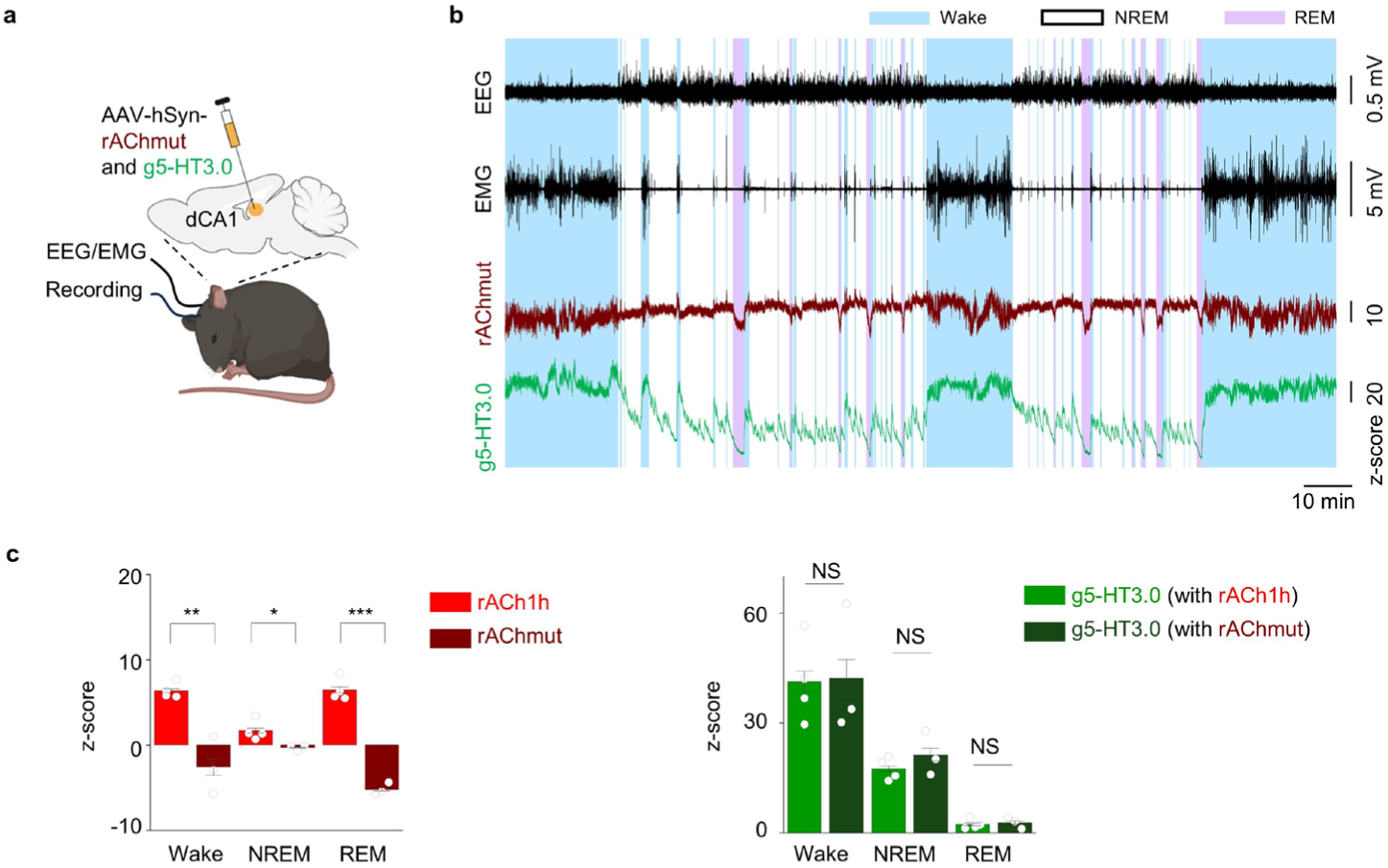
Representative rAChmut and g5-HT3.0 signals during the sleep-wake cycle in freely moving mice. **a**, Schematic illustration depicting the dual-color recording involving rAChmut and g5-HT3.0 during sleep–wake cycles for panel **b-c**. **b**, Representative traces of EEG, EMG, rAChmut (dark red) and g5-HT3.0 (green) during sleep–wake cycles in freely behaving mice. Bule shading, wake state; Pink shading, REM sleep. **c**, Group summary of rAChmut and g5-HT3.0 fluorescence in dCA1 compared to rACh1h during the wake state, NREM sleep, and REM sleep. The data of rACh1h and g5-HT3.0 (with rACh1h) is replotted from Fig.3c. mean ± s.e.m. n = 3 mice. Two-tailed Student’s t tests was performed. For rACh1h versus rAChmut, P = 0.003 during Wake, P = 0.037 between during NREM, P = 4.0×10^-5^ between during REM. NS, not significant.

**Extended Data Fig.7.**
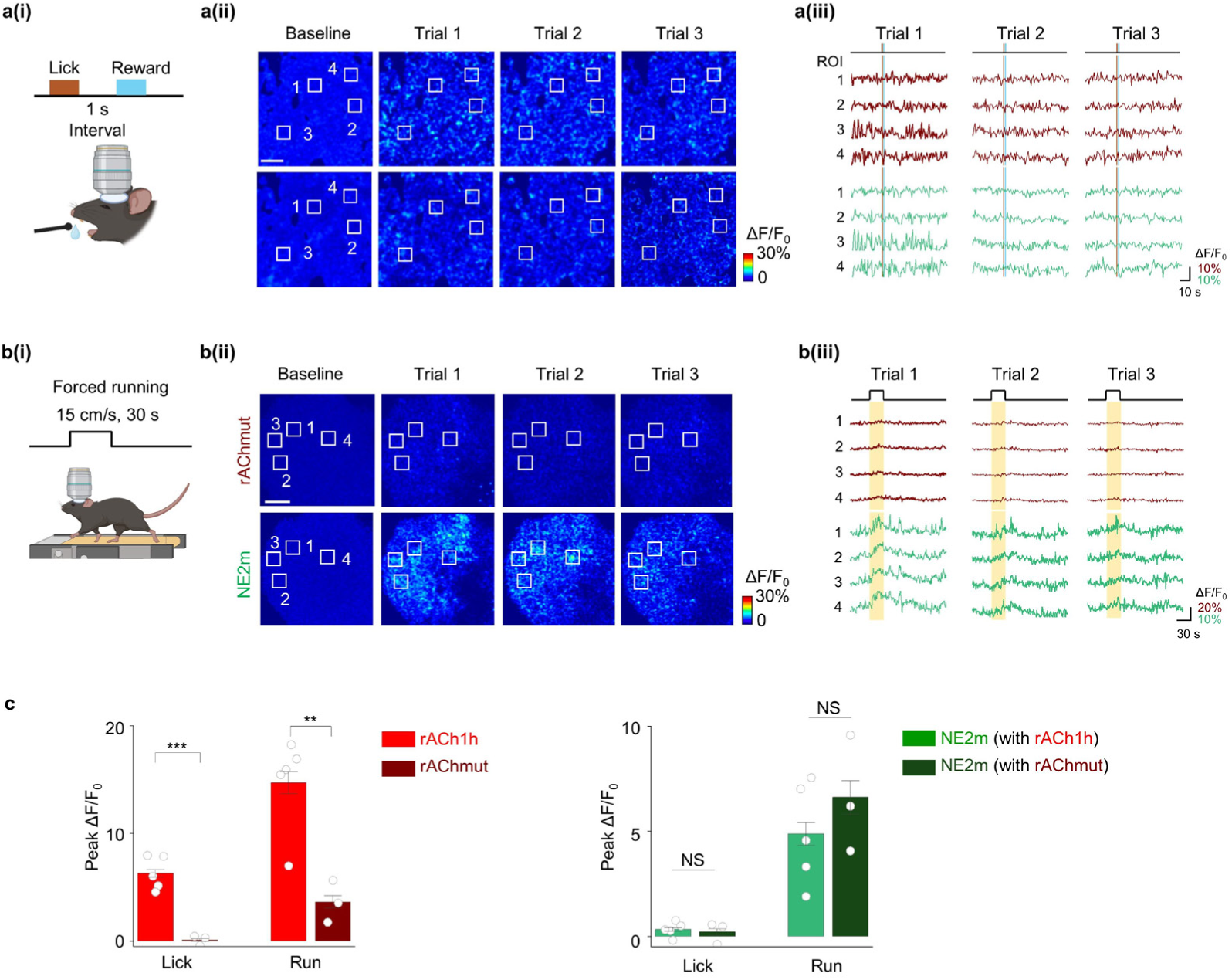
Representative rAChmut and NE signals in the cortex. **a**, Schematic cartoon illustrating water licking task (**a(i)**), representative response images (**a(ii)**) and typical traces (**a(iii)**) during three trials for rAChmut (Top) and NE2m (Bottom). Scale bar, 100 μm. **b**, Schematic cartoon illustrating forced running (**b(i)**), representative response images (**b(ii)**) and typical traces (**b(iii)**) during three trials for rAChmut (Top) and NE2m (Bottom). Scale bar, 100 μm. **c**, Group summary of rAChmut and NE2m peak response compared to rACh1h during the licking and running. The data of rACh1h and g5-HT3.0 (with rACh1h) is replotted from Fig.4h. mean ± s.e.m. n = 3 mice. Two-tailed Student’s t tests was performed. For rACh1h versus rAChmut, P = 6.0×10^-4^ in licking, P = 0.007 in running. NS, not significant.

